# Response metrics of T-cell activation models

**DOI:** 10.1101/2025.04.28.650960

**Authors:** Xabier Rey Barreiro, Jose Faro, Alejandro F. Villaverde

**Affiliations:** Universidade de Vigo, Department of Systems Engineering and Control, 36310 Vigo, Galicia, Spain; CITMAga, 15782 Santiago de Compostela, Galicia, Spain; Universidade de Vigo, Department of Biochemistry, Genetics and Immunology, 36310 Vigo, Galicia, Spain; Immunology Research Group, Galicia Sur Health Research Institute (IIS Galicia Sur), SERGAS-UVIGO, Spain

**Author notes:** Joint corresponding authors.

## Abstract

T-cell activation models describe the way in which antigens and T-cell receptors (TCRs) interact in order to trigger an immune response. A number of mathematical models of TCR-mediated T-cell activation have been proposed in the last decades. Often, the only measurable quantities available for the identification of these models are *E*_max_, the maximum achievable T cell activation level (over the background level), and *EC*_50_, that is, the antigen concentration that results in 50% of *E*_max_. In order to analyse model properties such as identifiability, observability, and sensitivity, it is necessary to know how *E*_max_ and/or *EC*_50_ relate to the model parameters. However, their expressions are only available for a few of the published models of T-cell activation. Here we present a general method to derive the expressions of *E*_max_ and/or *EC*_50_, and apply it to those models for which they are yet unknown. Our results expand the range of analyses that can be performed on T-cell activation models, contributing to enable their exploitation to understand and control the immune response.

## 1 Introduction

In vertebrates, the immune response consists of innate and adaptive strategies. The former is a first-line defense that reacts similarly to broad classes of pathogens, and may activate the latter one if a more targeted response is needed [1]. In contrast, the adaptive immune subsystem produces responses that are highly specific to each of the triggering antigens (Ag) [2]. Adaptive immune responses to protein Ags are based on interactions of T lymphocytes with Ag-presenting cells (APC) that lead to the activation of Ag-specific T cells. This lymphocyte activation is initiated by the binding of T-cell receptors (TCR) to their ligands, Ag-derived short peptides complexed with major histocompatibility complex proteins (pMHC) expressed on the membrane of APCs [2]. TCRs are membrane proteins composed of two chains, TCR*α* and TCR*β*, which can indirectly transduce a signal intracellularly through their tight, non covalent association with a membrane protein complex composed of two heterodimers and one homodimer, collectively called CD3 [2].

The detailed mechanisms through which this T-cell signalling eventually leads to cell fate decisions remain only partially known. Such limited mechanistic knowledge could be exploited, however, to support the development of methods to control the immune response. To this end, many mathematical models have been proposed to describe these T-cell activation processes in a quantitative way [3, 4, 5]. These dynamic models consist of sets of ordinary differential equations (ODEs), whose integration over time yields time-courses of the system variables [6]. The models are nonlinear, and contain parameters that are not directly measurable; therefore, they must be calibrated by optimizing the fit of the model output to experimental data [7]. Calibration leads to correct estimates only if the model is identifiable, i.e., if it is possible to determine the values of its parameters uniquely [8]. A related property, observability, informs about the possibility of inferring the unmeasured state variables from the model output [9, 10].

These structural properties of dynamic models depend on the model equations and on the definition of the output functions, i.e., of the variables —or functions thereof— that can be measured. Often, the experimentally measurable outputs consist of the maximum T-cell response level, *i*.*e*., the peak responses that can be obtained (*E*_max_), and the Ag concentration that induces a response that is halfway between the baseline and the maximum (*EC*_50_). Analysing these two outputs with respect to the aforementioned structural model properties requires mathematical expressions that relate these quantities to the model parameters and state variables. Furthermore, such expressions are also required for calculating the outputs’ sensitivity to changes in parameter values [11, 12]. However, these formulas are only available for a few of the models published to date [3]. While they can be obtained relatively straightforwardly in some cases, their derivation is in general non-trivial.

In this paper we address this problem in the following way. We begin by presenting, in Section 2, a general methodology for deriving expressions of *E*_max_ and *EC*_50_ for all models in which the response *R* increases monotonically with the amount of ligand, until reaching a plateau. Then, in Section 3 we provide those expressions for a comprehensive set of phenotypic T-cell activation models (*i*.*e*., models in which an output can be defined corresponding to the dynamics of a selected set of experimentally measurable quantities that determine a T-cell activation phenotype). This section also serves as a compendium of these models and their equations. For those models for which the expressions of *E*_max_ and *EC*_50_ had not been previously reported, their detailed derivations are given in the Appendix. Additionally, for one of the models —the serial triggering model described in subsection 3.13— we propose a modification of its equations, after discovering an important discrepancy between the conceptual description of the involved processes given in the original publication [13] and the equations presented therein. Lastly, we provide some concluding remarks in Section 4.

## 2 Methods

### 2.1 Phenotypic models of T-cell activation and parameters

The phenotypic models analysed in this work are summarized in Table 1, where references describing each model are included. Conceptual diagrams are shown in Fig. 1. All models have a minimum set of common parameters (most of which are kinetic rate constants). Both common and specific parameters are indicated in Fig. 1.

**Table 1:**
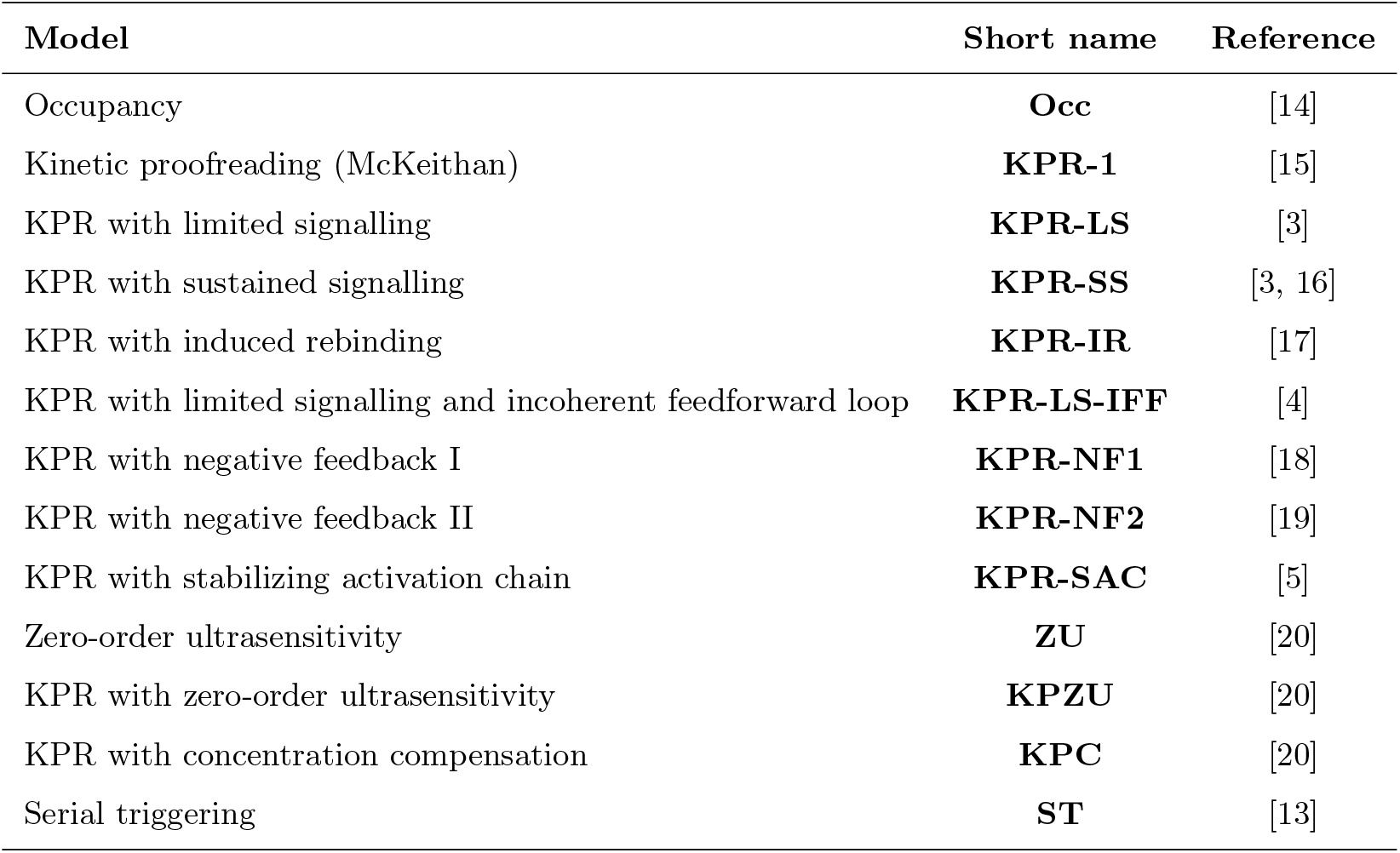
Phenotypic models analysed in this work.

**Figure 1:**
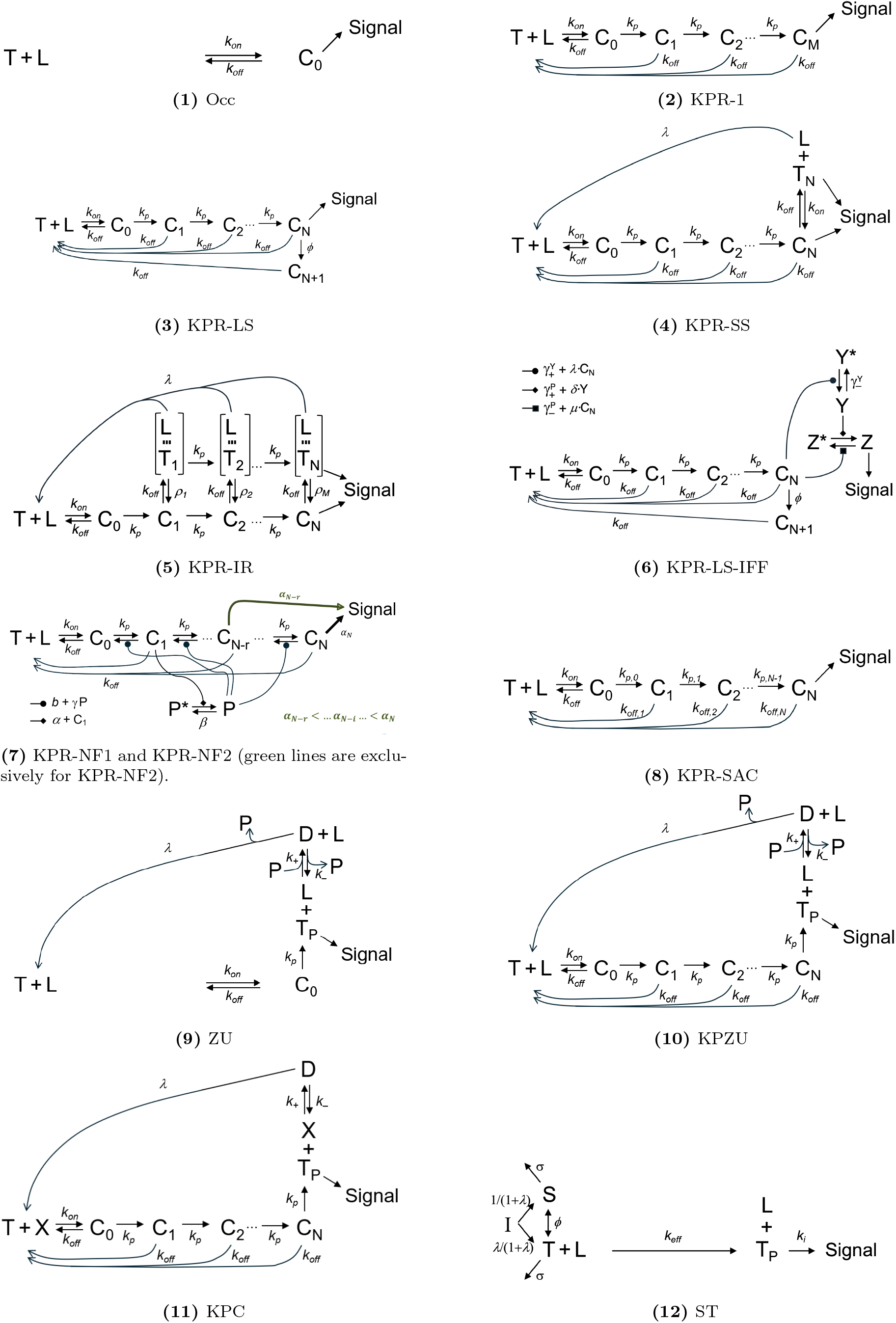
Diagrams describing the different models analysed in this paper.

### 2.2 Modeling framework and notation

T-cell activation models are typically described by a system of ordinary differential equations (ODEs) that can be written in the following general form:

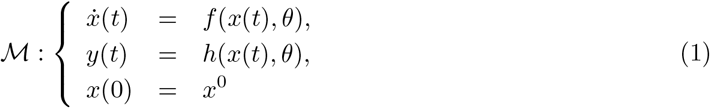

where *θ* = (*θ*_1_, …, *θ*_*p*_) is the parameter vector; *x*(*t*) = (*x*_1_(*t*), …, *x*_*n*_(*t*)) is the state vector with initial conditions given by *x*(0) = (*x*_1_(0), …, *x*_*n*_(0)); *y*(*t*) = (*y*_1_(*t*), …, *y*_*m*_(*t*)) is the output vector; and *f* (*x*(*t*), *θ*) and *h*(*x*(*t*), *θ*) are non-linear functions. We will often simplify the notation by dropping the dependence on *t*. All vectors are real-valued, and typically have positive values.

The ODEs defining each phenotypic model are presented in Section 3 with a brief description of the corresponding model.

### 2.3 Calculation of *E*_max_ and *EC*_50_

The phenotypic models considered in this work describe distinct processes, initiated by TCR-pMHC binding, resulting in the activation of TCRs (denoted also triggered TCRs) and the subsequent intracellular T cell signalling. In order to establish in each model a connection between total ligand per APC and T cell activation, we followed the approach in [3] and considered the outputs *E*_max_ and *EC*_50_ as two measures of T cell activation.

In this section we describe a procedure to derive, for all phenotypic models, the corresponding expressions for *E*_max_ and *EC*_50_ as functions of parameters and total ligand and TCR per cell. The main steps of this procedure are described in the following subsection.

#### 2.3.1 Deriving the expressions of *E*_max_ and *EC*_50_ in T-cell activation ODE models

**Step 1:** *Steady-state*. First, assume that the system is in steady-state and, therefore, set to zero the differential equations given by *f* (*x*(*t*), *θ*). Also, consider the following conservation equations, respectively, for the ligands and receptors:

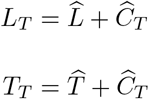

where *L*_*T*_, *T*_*T*_ and 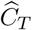 are, respectively, the total cell density of pMHC, TCR and TCR-pMHC complexes, and 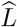 and 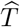 are, respectively, the cell density of unbound pMHC and TCR at steady-state.

**Step 2:** *Define* 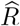, *the T-cell response level at steady-state, in terms of triggered TCRs, and obtain an expression for* 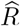 *as a function of total ligand, L*_*T*_. That expression can be obtained via two approaches:

- (2A): By incorporating the conservation equations within the equations that model the dynamics of free receptors and ligands, the cell density of free ligands and TCRs can be expressed in terms of 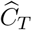. Subsequently, using the remaining equations in the system at steady-state, derive an expression linking 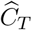 to *R*.
- (2B): If the equations that describe the evolution of free receptors and ligands contain some state variables in addition to 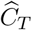, it may be necessary to find a direct relationship between 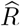 and *L*_*T*_ through each model equation.

**Step 3:** *Obtain an expression for E*_*max*_, *the maximal response level*. For all models but four (KPR-LS-IFF, KPR-NF1, KPR-NF2, and KPC, considered below as special cases), the corresponding value of 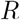 grows monotonically with *L*_*T*_, reaching a maximal level or plateau at ligand saturation. Following [3] we define *E*_max_ as the value of 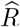 in the limit of excess ligand or ligand saturation, that is, in all these models 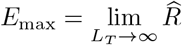.

To that end, in the first approach (2A), compute first the limit of 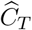 for *L*_*T*_ *→ ∞*, and then substitute this result into the expression for *R*. In the second approach (2B), the limit *L*_*T*_ *→ ∞* is directly computed after non-dimensionalizing the variables.

**Step 4:** *Obtain EC*_50_. Impose the condition 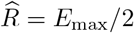 and substitute the expressions obtained in steps 2 and 3 for 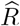 and *E*_max_. This yields an expression in terms of *L*_*T*_, *T*_*T*_, and all or a subset of the system parameters. Then solve this expression for *L*_*T*_ in terms of *T*_*T*_ and parameters. Thus, *EC*_50_ = *L*_*T*_ when *R* = *E*_max_*/*2.

#### 2.3.2 Illustration of the methodology

Here we apply the above procedure to McKeithan’s kinetic proofreading (KPR) model [15, 3] described below in subsection 3.2. Its corresponding ODE system is defined in that subsection by Eqns. (19). In this model the T-cell activation level is given by:

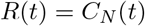

**Step 1:** As indicated above, we denote the system state variables at steady-state with a hat accent over the variable’s name. Then the model equations at steady-state are:

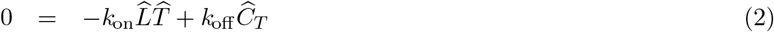

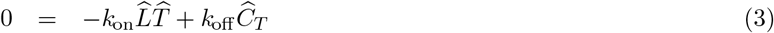

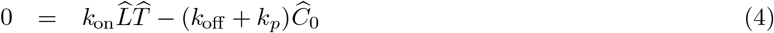

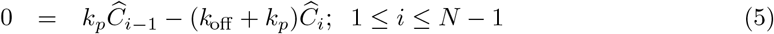

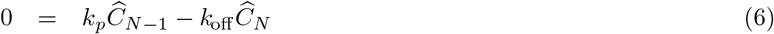

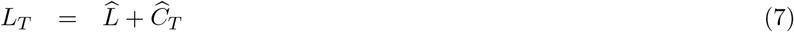

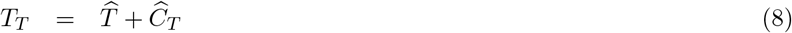

**Step 2:** From Eqn. (5) one has 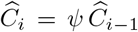, with 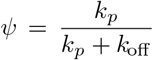, and applying this equation recursively, one obtains:

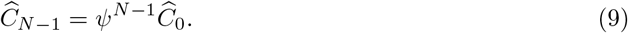

Taking into account that *k*_off_ */k*_*p*_ = (1 *− ψ*)*/ψ*, and inserting Eqn. (9) into Eqn. (6) yields 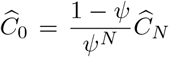. Plugging now this equation into the equation that defines the total number of ligand-receptor complexes yields:

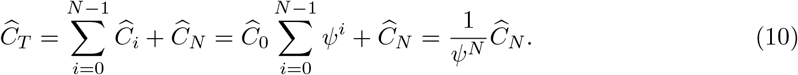

Therefore:

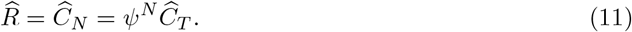

Now, from Eqn. (2) one has:

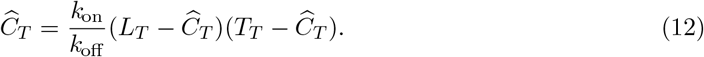

Solving this equation for 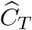 gives:

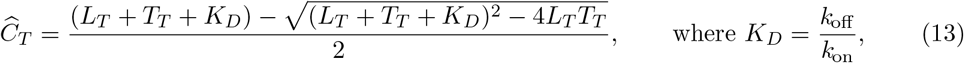

and inserting this result into Eqn. (11) provides an expression for *R* in terms of *L*_*T*_.

**Step 3:** Taking into account Eqn. (11), calculating 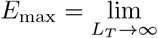 *R* amounts to calculating 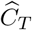 for *L*_*T*_ *→ ∞*. In order to do that, it suffices to reorganise Eqn. (12) and take the limit *L*_*T*_ *→ ∞*:

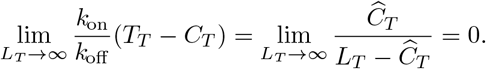

Therefore, 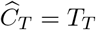 in the limit *L*_*T*_ *→ ∞*. Hence:

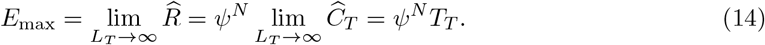

**Step 4:** Let *R*_50_, 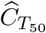, and 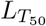 denote, respectively, the half-maximal response, the total complexes at *R*_50_, and the total ligand dose that induces the *R*_50_ response. Then, from Eqns. (11) and (14) one has:

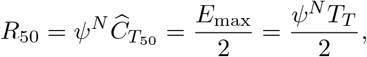

and, therefore,

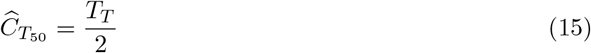

Now, from Eqn. (13) and using Eqn. (15), one has at 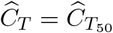:

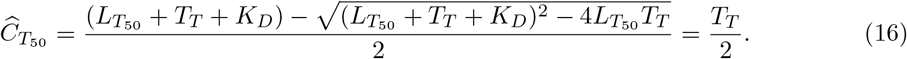

Finally, solving Eqn. (16) for 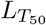 yields:

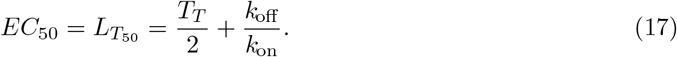

Figure 2 shows a graphical summary of these results for a particular set of parameter values.

**Figure 2:**
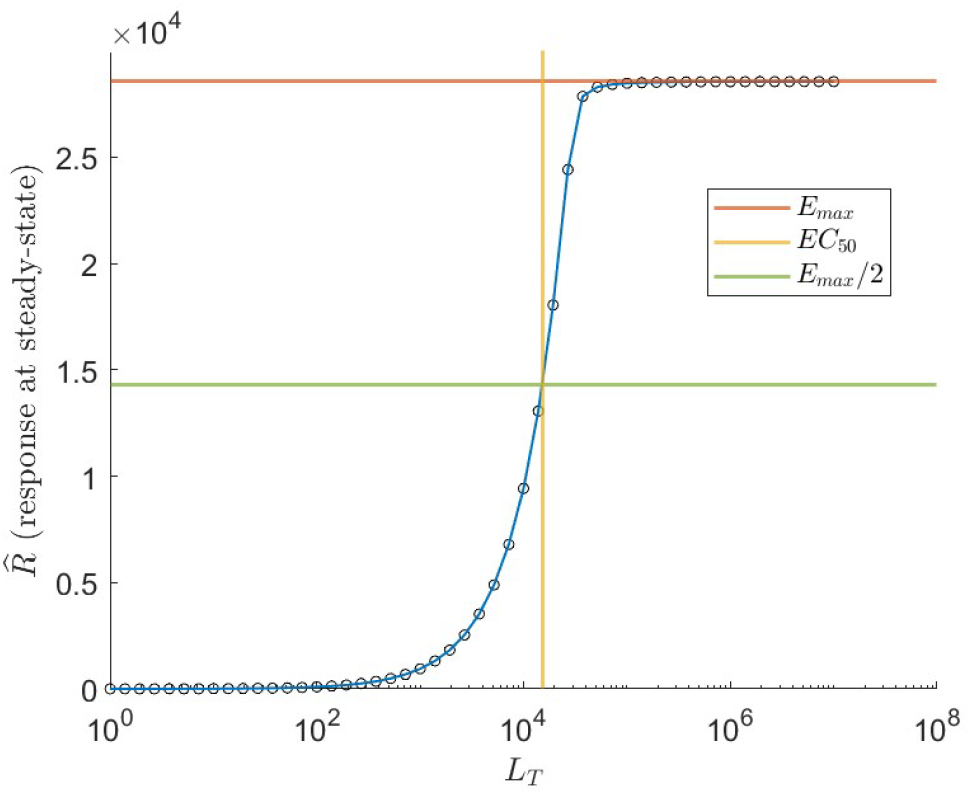
*KPR-1* model. The response at steady-state, 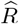, as a function of total ligand. Black circles correspond to values of *R*(*t*) obtained with the ODE system at *t* = 300 *s*. The horizontal red line shows the value of *E*_max_ computed with Eqn. (14), while the green line corresponds to *E*_max_*/*2. The vertical yellow line shows the value of *EC*_50_ obtained with Eqn. (17). Parameter values: *k*_*on*_ = 10^*−*5^ *s*^*−*1^[]^*−*1^, *k*_off_ = 0.01 *s*^*−*1^, *k*_*p*_ = 1 *s*^*−*1^, and *T*_*T*_ = 3 *×* 10^4^ TCRs/cell. The symbol [] indicates a concentration unit (number of molecules per cell) for membrane and cytoplasmic molecules.

## 3 Results: calculation of 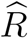, *E*_**max**_ and *EC*_50_ for T-cell phenotypic models

In this section we briefly present the currently existing phenotypic models of T-cell activation (Table 1) with their defining equations, in approximate order of increasing complexity. For most of them we provide the corresponding expressions for 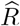, *E*_max_ and *EC*_50_. We start with those few models for which these expressions were already available in the literature [3] (Sections 3.1 to 3.3). For them, we give the published results without further calculations. In section 3.4 we deal with the KPR with sustained signalling model. While this model was also analysed in reference [3], in that publication only an implicit-form expression was provided for 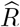 and *E*_max_. Therefore, we reanalysed here this model, and in that section we provide the corresponding closed-form expressions for those two quantities in terms of *L*_*T*_ and *T*_*T*_. For most of the remaining models (Sections 3.5 to 3.13) we give, for the first time, the corresponding expressions for *R, E*_max_ and *EC*_50_. Since in general their derivation departs to varying degrees from the procedure outlined in subsection 2.3.2, we provide their detailed calculations in the Appendix.

To highlight the similarities and differences among the models, we show their conceptual schemes in Fig. 1 using a similar layout for all of them. The code and calculations used to produce the results can be accessed at the following repository: https://github.com/Xabo-RB/Inmunology-analysis.git.

### 3.1 Occupancy model (*Occ*)

The occupancy model presented in [14] assumes that T-cell activation is proportional to the number of TCRs occupied by pMHC complexes, without further delays. Its equations are given by:

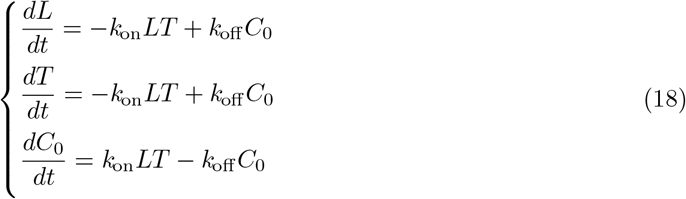

The response or signalling state in this model is defined as *R*(*t*) = *C*_0_(*t*). For this model, the expressions for 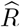, *E*_max_ and *EC*_50_ can be found in the literature [3], and are as follows:

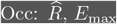, and *EC*_50_.

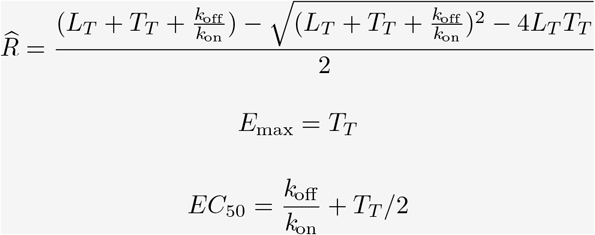

### 3.2 KPR McKeithan (*KPR-1*)

This is the first and simplest model of lymphocyte receptor activation through kinetic proofreading (KPR) proposed originally by McKeithan [15]. According to this mechanism, interaction between the TCR and the peptide-MHC complex does not immediately lead to signalling; instead, TCRs in complexes with pMHC ligands must undergo *N* sequential modifications (phosphorylation of the CD3 cytoplasmic tails) before triggering an effective immune response. In the KPR-1 model the response is defined as *R*(*t*) = *C*_*N*_ (*t*). The ODE system defining this model is the following:

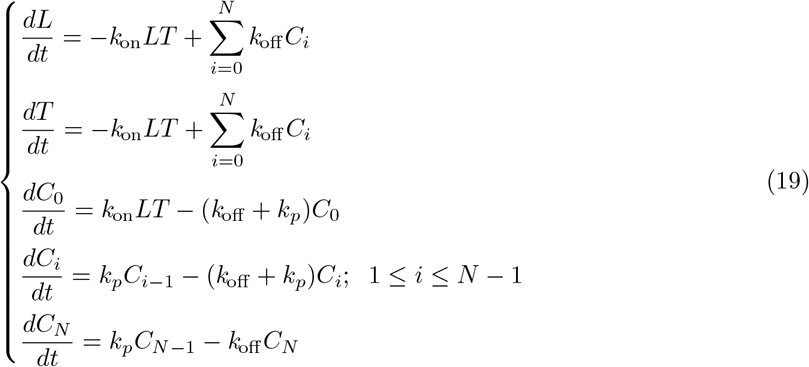

and the corresponding 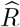, *E*_max_ and *EC*_50_ are:

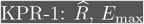, and *EC*_50_.

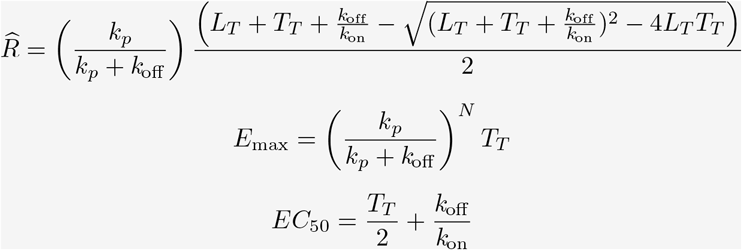

### 3.3 KPR limited signalling (*KPR-LS*)

This extension of the KPR-1 model, proposed in [3], assumes that T-cell receptors signal for a limited time after reaching an activated state. For this model, *R*(*t*) = *C*_*N*_ (*t*). This model is defined by the following ODE system:

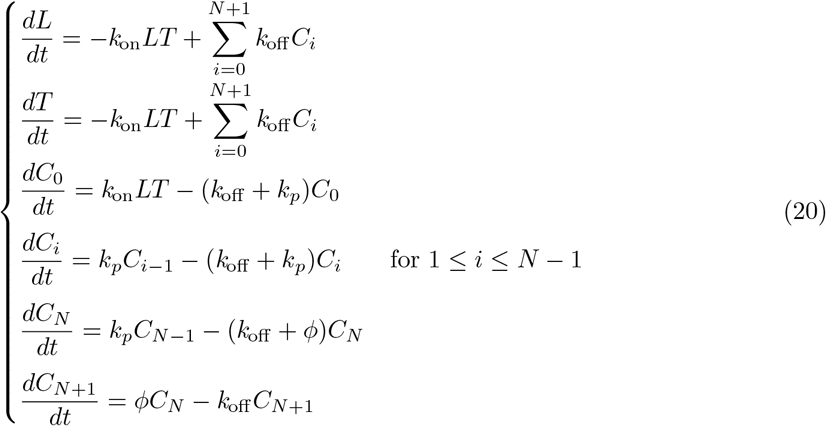

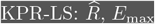 and *EC*_50_.

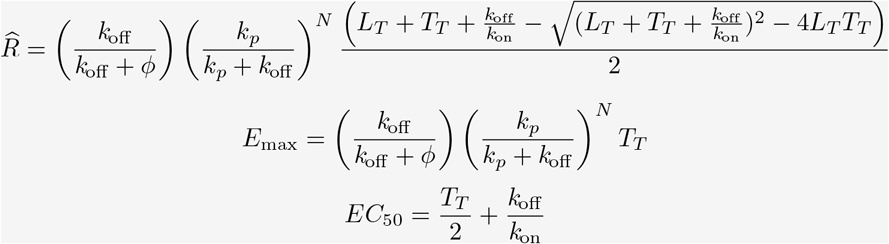

### 3.4 KPR sustained signalling (*KPR-SS*)

This model [16] combines kinetic proofreading and serial engagement mechanisms, and defines the response as *R*(*t*) = *C*_*N*_ (*t*) + *T*_*N*_ (*t*). Its ODE equations are:

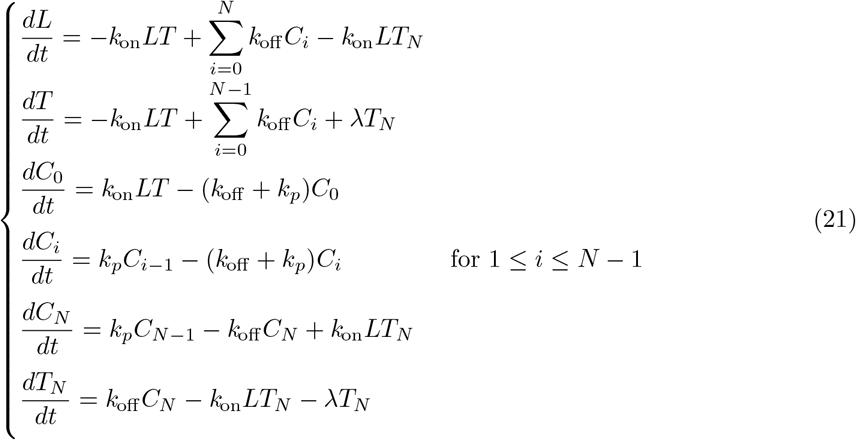

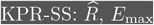 and *EC*_50_.

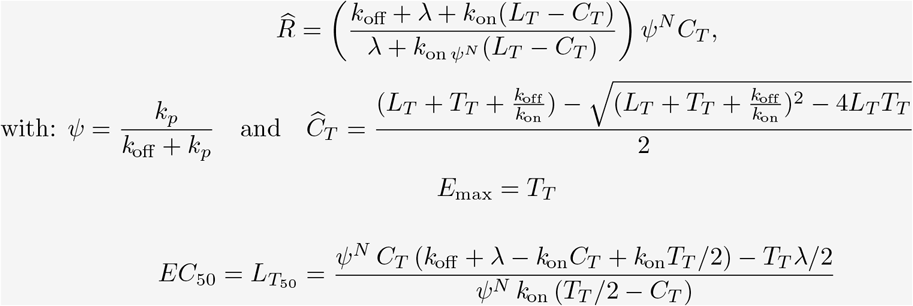

Note that the expression for *EC*_50_ or 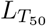 written above, which is taken from [3], depends implicitly on *L*_*T*_ through *C*_*T*_. Therefore, given in this form, the equation for *EC*_50_ must be numerically solved for *L*_*T*_. To solve this issue, we performed the corresponding calculations and derived the following closed-form expression for *EC*_50_ previously unavailable in the literature (see Appendix, subsection **KPR sustained signalling (KPR-SS)**):

KPR-SS: *EC*_50_.

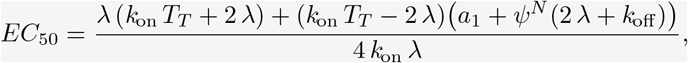

with,

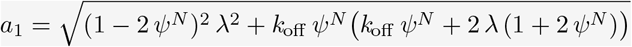

### 3.5 KPR with induced rebinding (KPR-IR)

The model of *KPR with induced rebinding* assumes some changes in the receptor that could induce the rebinding to a pMHC [17]. Here we characterise this possible rebinding before fully dissociating by considering an intermediate ligand-receptor state denoted 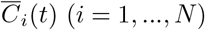. These are intermediate [*T*_*i*_-*L*] encounter complexes characterised by *T*_*i*_ and *L* being within the reaction distance but not bound [21]. The response is defined in this model as 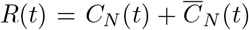, where 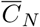 denotes the intermediate complex state [*T*_*N*_ -*L*]. The ODE system defining this model is:

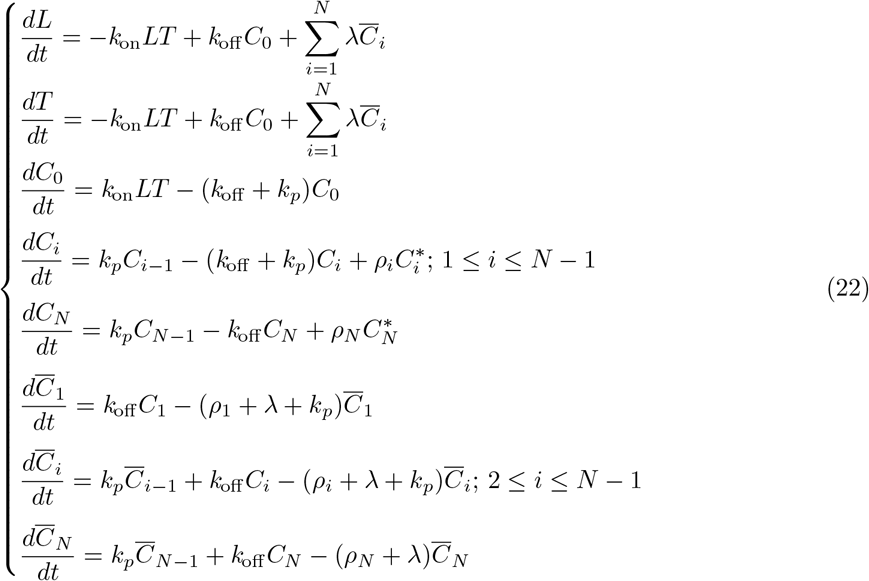

In order to derive an approximate version of this ODE system that could be solved analytically, we introduce the following hypothesis: due to experimental limitations, it is not possible to discriminate between the two molecular states *C*_*i*_ and 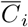. In particular, when re-binding after unbinding occurs fast enough, we cannot distinguish between unbound but encounter-oriented states and bound states [21]. In mathematical terms, this means that, in a coarse-grained observation timescale, we can safely assume that 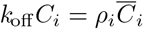 (in statistical thermodynamics this hypothesis is often referred to as local steady state hypothesis). Then, defining 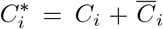, one has: 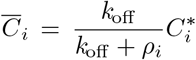 and 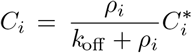. Consequently, the above ODE system reduces to the following one:

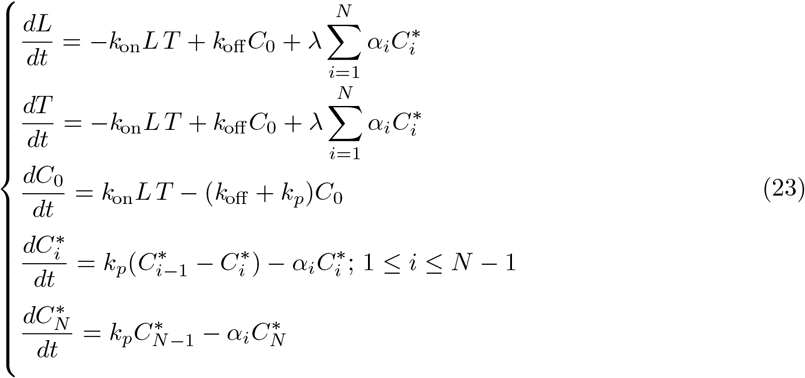

where 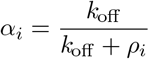. Note that in this approximate system 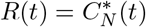.

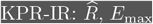 and *EC*_50_.

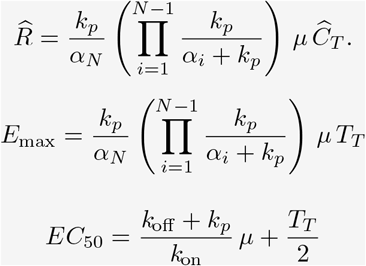

Here the auxiliary parameter *µ* and state variable 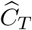 are as defined, respectively, in Eqns. (40) and (42) in the Appendix.

### 3.6 KPR limited signalling and incoherent feed-forward loop (*KPR-LS-IFF*)

An incoherent feed-forward loop is a signalling mechanism where an input activates the target signal through one route, and inhibits it through another one. In the KPR-LS-IFF, such a circuit is combined with the limited signalling mechanism described above [22, 4]. The response is the last state variable, *R*(*t*) = *Z*(*t*). This model is defined by the following ODE system:

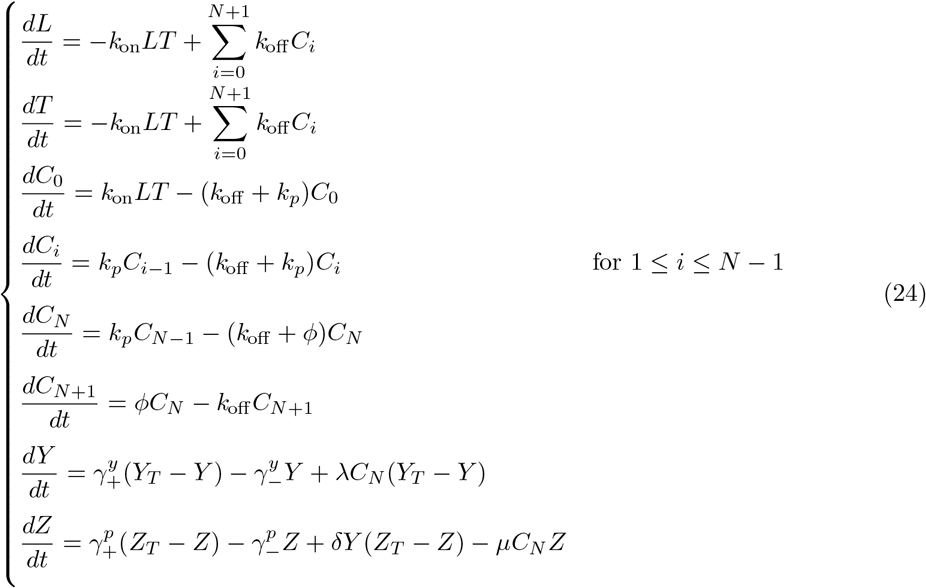

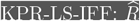.

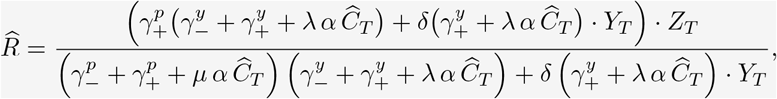

with,

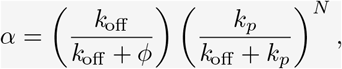

and

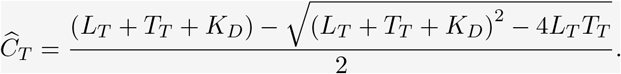

In this model, the response 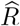 as a function of *L*_*T*_ displays, for many reasonable parameter values, a bell-shaped behaviour (see Fig. 3). Therefore, to determine *E*_max_ it is necessary to compute first the derivative 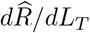, and subsequently solve the equation 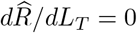 to identify the corresponding value of *L*_*T*_. Denoting this specific value as 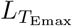, the expression for *E*_max_ is given by 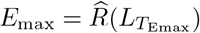. Owing to their big size, the expressions for both, *E*_max_ and *EC*_50_ are only available in our online repository (https://github.com/Xabo-RB/Inmunology-analysis.git).

**Figure 3:**
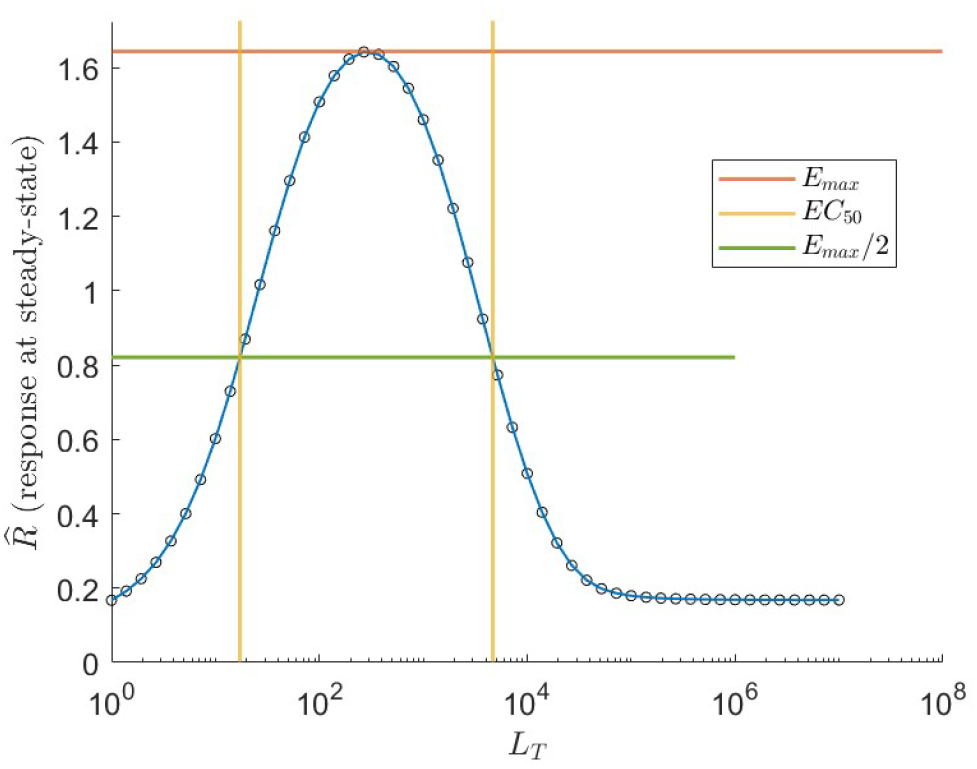
*KPR-LS-IFF* model. The response at steady-state, 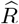, as a function of total ligand shows a bell-shape behaviour for many biologically reasonable parameter values. The blue line was obtained using the general equation for 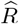. Black circles correspond to values of *R*(*t*) obtained by solving the ODE system at *t* = 300 *s* for a large range of *L*_*T*_ values. The horizontal red and green lines show, respectively, the computed values of *E*_max_ and *E*_max_*/*2. The vertical yellow lines correspond to the two possible values of *EC*_50_. Parameter values: *n* = 5 kinetic steps, *k*_on_ = 10^*−*5^ *s*^*−*1^[]^*−*1^, *k*_off_ = 0.05 *s, k*_*p*_ = 0.04 *s, ϕ* = 0.04 *s, γ*_+_ = 1 *s*^*−*1^, *γ*_*−*_ = 500 *s*^*−*1^, *λ* = 100 *s*^*−*1^[]^*−*1^, *δ*= 100 *s*^*−*1^[]^*−*1^, *µ* = 500 *s*^*−*1^[]^*−*1^, *Y*_*T*_ = 100 molec/cell, *Z*_*T*_ = 2.5 molec/cell, and *T*_*T*_ = 3 *×* 10^4^ TCRs/cell. The symbol [] indicates a concentration unit (number of molecules per cell) for membrane and cytoplasmic molecules.

### 3.7 KPR with negative feedback 1 (*KPR-NF1*)

This model extends the basic KPR mechanism by adding a negative feedback loop mediated by Src homology 2 domain phosphatase-1 (SHP-1) [18, 23]. The response is defined here as *R*(*t*) = *C*_*N*_ (*t*), and the corresponding ODE system is:

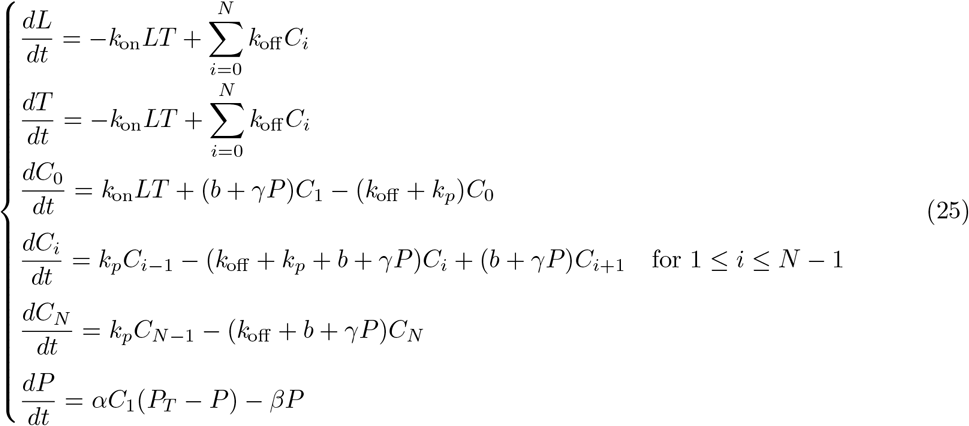

As mentioned above and briefly discussed in [3, 4, 18], the *KPR-NF1* exhibits, for many parameter combinations and *N >* 2 kinetic steps, a bell-shaped behaviour in the relationship between the response 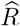 and the total amount of ligands presented, *L*_*T*_. This characteristic behaviour is shown, for *N* = 5, in panel (2) of Figure 4. For *N* = 2, the *E*_max_ value is attained when *L*_*T*_ approaches infinity, as in the other models. The expression for *R* cannot be shown here due to its big size; we provide it at: https://github.com/Xabo-RB/Inmunology-analysis.git.

**Figure 4:**
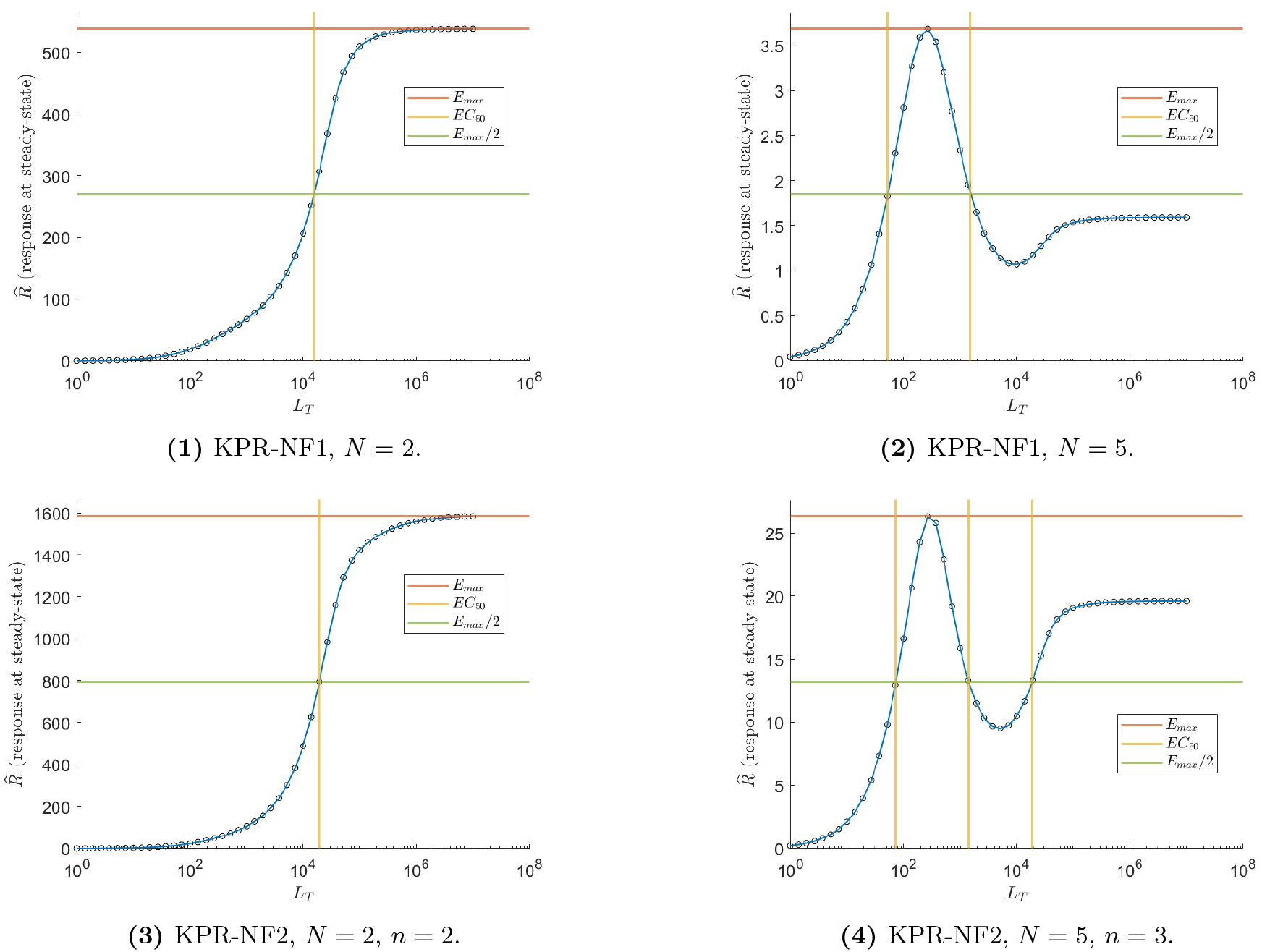
*KPR-NF1* and *KPR-NF2* models. Response level, *R*, as a function of *L*_*T*_, for the KPR-NF1 model (first row) and the KPR-NF2 model (second row). Left panels, results for two kinetic proofreading steps (*N* = 2) for both models, and *n* = 2 for KPR-NF2. Right panels, results for *N* = 5 for both models, and *n* = 3 for KPR-NF2. Parameter values: *α/β* = 0.002, *b* = 0.04 *s*^*−*1^, *γ* = 1.0 *×* 10^*−*6^ *s*^*−*1^[]^*−*1^, *k*_on_ = 1.0 *×* 10^*−*5^ *s*^*−*1^[]^*−*1^, *k*_off_ = 0.05 *s*^*−*1^, *k*_*p*_ = 0.09 *s*^*−*1^, *T*_*T*_ = 3 *×* 10^4^ TCRs/cell, and *S*_*T*_ = 6 *×* 10^5^ phosphatase molecules/cell. In panel (4) *k*_*p*_ = 0.13 and *k*_off_ = 0.03. The symbol [] indicates a concentration unit (number of molecules/cell) for membrane and cytoplasmic molecules.

For *N >* 2, the *E*_max_ value must be obtained by analysing the derivative of an expression that describes, under steady-state conditions, the relationship between 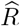 and *L*_*T*_. For *N* = 5 and a standard set of parameter values (see Table 2 in the Appendix) we obtained the derivative numerically, using Mathematica [24]. In this case, the response *R* as function of *L*_*T*_ has five critical points, two of which are complex solutions and one is negative. Therefore, only two real positive possible solutions remain for the maximal response. Since *E*_max_ is, by definition, the solution with the largest numerical value, it corresponds to the local maximum of the curve shown in Fig. 4, panel (2). Thus, unlike most phenotypic models of TCR activation, in many reasonable parameter regimes the KPR-NF1 model does not reach *E*_max_ at ligand saturation but at relatively low ligand density in APCs. A limited analysis of the parameter impact on the behaviour of *R vs L*_*T*_ showed that the parameters that affect most this behaviour are *k*_off_, *k*_*p*_ and the ratio *α/β*. Because the behaviour of the response *R* as a function of *L*_*T*_ has in many reasonable parameter regimes a bell shape with a rebound to a plateau, the ligand dose that produces a half-maximal T cell response, *i*.*e*. 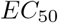, may have up to three different positive solutions. In the present example (Fig. 4, panel (2)) there are two solutions: 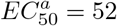 and 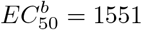 ligands per APC.

**Table 2:** Parameter values used in the analyses.

| Parameter | Value | Description | Ref. |
| --- | --- | --- | --- |
| $k_{\text{on}}$ | $5.0 \times 10^{-5}$ | Binding rate between TCR and ligand | [4] |
| $k_{\text{off}}$ | $1.0 \times 10^{-2}$ | Unbinding rate between TCR and ligand | [4] |
| $T_T$ | $3.0 \times 10^4$ | Number of TCRs | [4] |
| $k_p$ | $1.0 \times 10^0$ | Forward rate in proofreading | [4] |
| $P_T$ | $6.0 \times 10^5$ | Total number of SHP-1 | [5] |
| $\beta/\alpha$ | $5.0 \times 10^2$ | SHP-1 deactivation-activation ratio | [5] |
| $b$ | $4.0 \times 10^{-2}$ | Spontaneous dephosphorylation rate by SHP-1 | [2] |
| $\gamma$ | $1 \times 10^{-6}$ | Dephosphorylation rate by SHP-1 | [5] |
| $\gamma$ | $5.0 \times 10^2$ | Ratio of background inhibition and activation (*KPR-LS-IFF*) | [7] |
| $\phi$ | $9.0 \times 10^{-2}$ | Forward rate from signalling to non-signalling | [8] |
| $\rho$ | $1.0 \times 10^3$ | Rebinding rate between TCR and ligand | [1] |
| $r$ | $1.5 \times 10^0$ | Factor for increasing dephosphorylation rate | [3] |
| $r_p$ | $1.03 \times 10^0$ | Factor for increasing forward rate | [3] |
| $Y_T$ | $1.0 \times 10^2$ | Total number of Y nodes | [7] |
| $P_T^*$ | $2.5 \times 10^0$ | Total number of P nodes | [7] |
| $\delta$ | $1.0 \times 10^2$ | Activation parameter | [7] |
| $\mu$ | $5.0 \times 10^2$ | Inhibition parameter | [7] |
| $\lambda$ | $1.0 \times 10^2$ | Amplification parameter for *KPR-LS-IFF* | [7] |
| $\lambda$ | $4.0 \times 10^{-2}$ | Reaction constant | [6] |
| $k_+$ | $1.0 \times 10^{-1}$ | Reaction constant | [6] |
| $k_-$ | $5.0 \times 10^{-2}$ | Reaction constant | [6] |

### 3.8 KPR with negative feedback 2 (*KPR-NF2*)

The KPR-NF1 model was later relaxed [19] with respect to the definition of the response *R*. Nevertheless, the ODE system defining this model is still the same as that of the KPR-NF1 model and, therefore, all calculations but *R* remain in this model equal to the corresponding ones in the KPR-NF1 model. The KPR-NF2 model departs from the previous one by hypothesizing that not only TCRs in the *C*_*N*_ complexes contribute to T cell signalling, but also TCRs within the final *n* complexes of the proofreading chain (*i*.*e*., from *C*_*N−n*+1_ to *C*_*N*_), although with a contributing weight increasing from *N − n* + 1 to *N*. To account for this modification we define the following response, which assigns increasing weights *α*_*i*_ to the contributions of each *C*_*i*_:

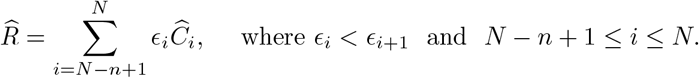

The second row of Figure 4 illustrates the same effect observed in the *KPR-NF1* model: for *N >* 2 and *n >* 1 the peak response does not occur at ligand saturation (*i*.*e*., for *L*_*T*_ *→ ∞*), but at relatively low ligand levels. Similarly to the KPR-NF1 model, depending on parameter values, particularly those of *k*_off_, *k*_*p*_ and the ratio *α/β*, there may be up to three different and biologically relevant solutions for *EC*_50_. As in the previous model, the code for 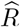 is available in our repository for *N* = 2 and *n* = 2 (https://github.com/Xabo-RB/Inmunology-analysis.git).

### 3.9 KPR with stabilizing activation chain (*KPR-SAC*)

The “Stabilizing Activation Chain” mechanism [5] relaxes the assumptions that all phosphorylation steps in the KPR mechanism (*i*.*e*., *C*_*i*_ *→ C*_*i*+1_) occur with the same rate constant *k*_*p*_, and that all unbinding reactions (*i*.*e*., *C*_*i*_ *→ L* + *T*) have the same rate constant *k*_off_. This model was designed to lead to different responses for foreign and self-peptides. Thus, in this model it is assumed, on the one hand, that the phosphorylation steps *C*_*i*_ *→ C*_*i*+1_ take place with rate constant 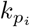, with 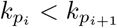, and, on the other hand, that the unbinding reactions *C*_*i*_ *→ L* + *T* have rate constants *k*_off *i*_, with *k*_off *i*_ *> k*_off *i*+1_. In this model the response is defined as *R*(*t*) = *C*_*N*_ (*t*), and the defining ODE system is:

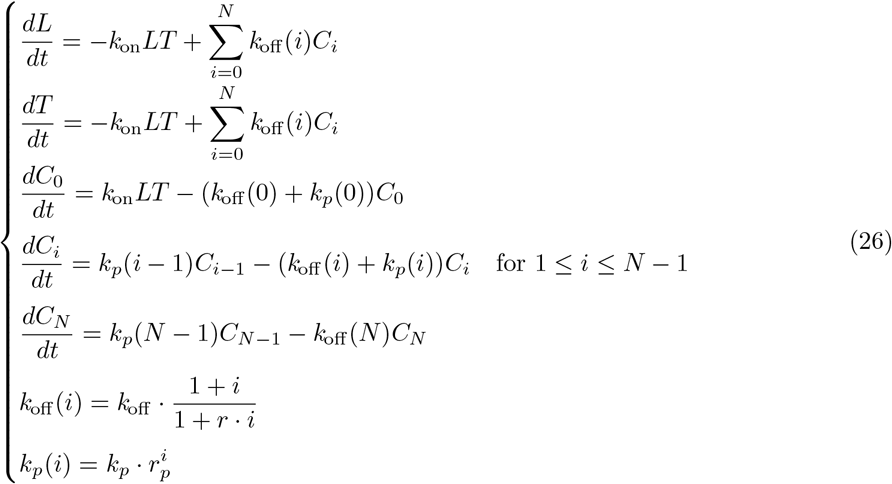

where *r* and *r*_*p*_ are constants.

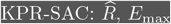 and *EC*_50_.

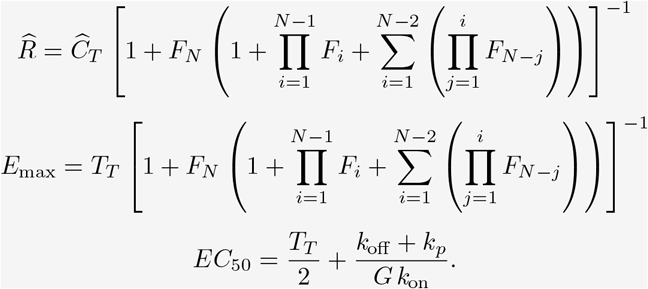

Expressions for *F*_*i*_, *F*_*N*_, 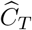, and *G* are given, respectively, in Eqns. (58), (59), (66) and (69) in the Appendix.

### 3.10 Zero-order ultrasensitivity model (*ZU*)

Zero-order ultrasensitivity [25] is a mechanism that enhances sensitivity to changes in ligand concentration and leads to fast responses, at the expense of decreasing specificity and the ability to discriminate foreign *vs* self-antigens [20]. In this model, *R*(*t*) = *T*_*p*_(*t*), and the ODE system defining this model is the following:

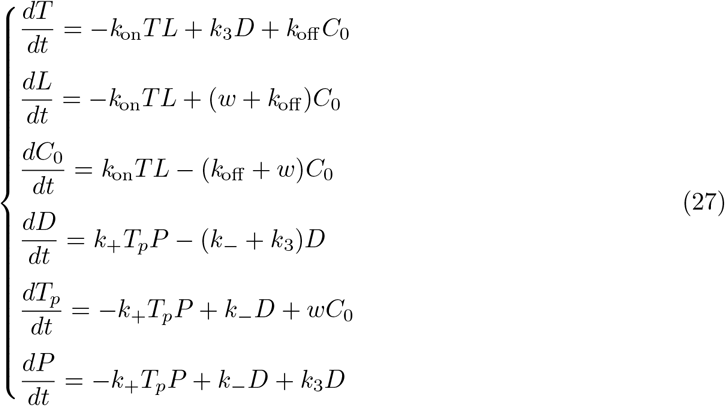

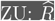 and *E*_max_.

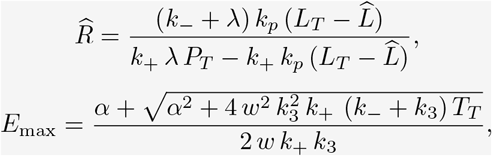

where:

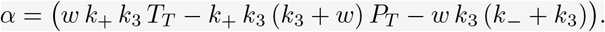

An expression for 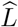 is given by a solution to Eqn. (77) in the Appendix. The formula for *EC*_50_ is exceedingly large to be shown here; instead, we provide it in our online repository (https://github.com/Xabo-RB/Inmunology-analysis.git).

### 3.11 KPR with zero-order ultraspecificity (*KPZU*)

The KPR with zero-order ultraspecificity model combines KPR and zero-order ultrasensitivity [20]. In this way it can balance specificity, sensitivity, and speed. The response is given by *R*(*t*) = *T*_*p*_(*t*), and the equations defining this model are the following:

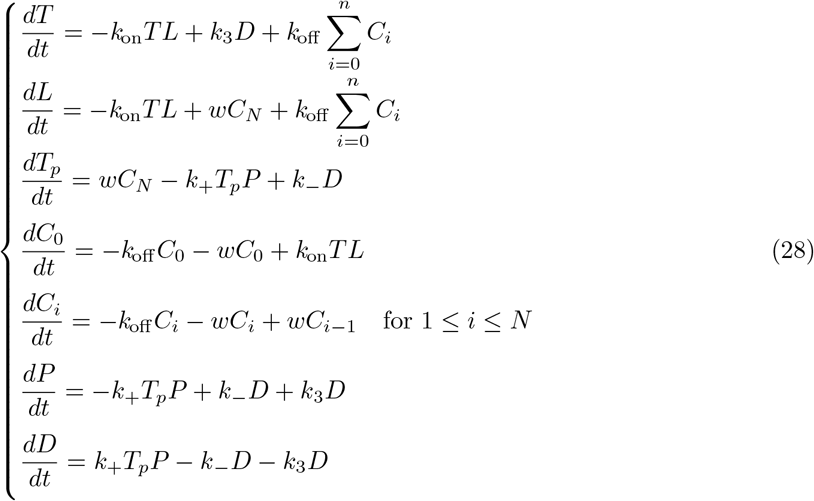

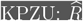 and *E*_max_.

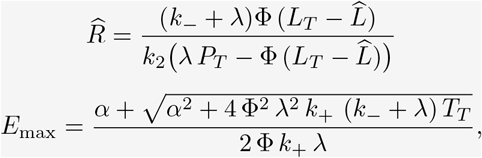

where:

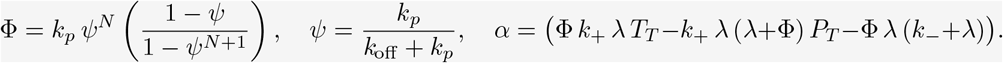

As was the case for the *ZU* model, an expression for 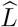 is given by the solution of Eqn. (81) in the Appendix, and a general formula for *EC*_50_ is provided in our online repository (https://github.com/Xabo-RB/Inmunology-analysis.git).

### 3.12 KPR with concentration compensation (*KPC*)

The *KPC* model modifies the *KPZU* model by assuming a mechanism where the same molecule controls receptor phosphorylation and dephosphorylation [20]. Thus, the system can exhibit low sensitivity to high concentrations of non-target ligands and high sensitivity to low concentrations of target ligands. As in the previous model, *R*(*t*) = *T*_*p*_(*t*). The ODE system defining this model is:

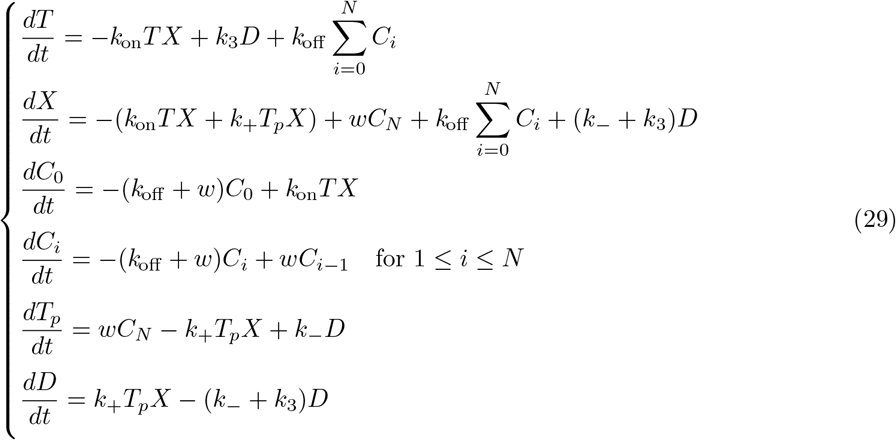

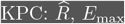 and *EC*_50_.

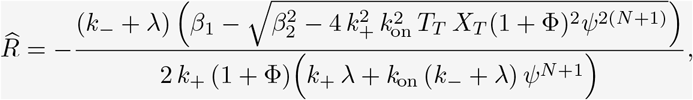

where:

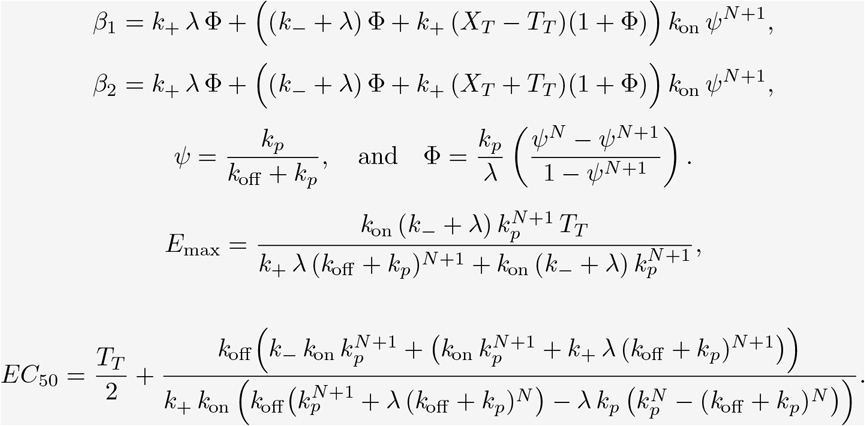

This is an atypical model in that the response at steady-state, 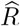, as a function of total ligand shows an inverted sigmoidal behaviour (Fig. 5), with a maximum at *L*_*T*_ *→* 0 and decreasing to 0 for *L*_*T*_ *→ ∞*.

**Figure 5:**
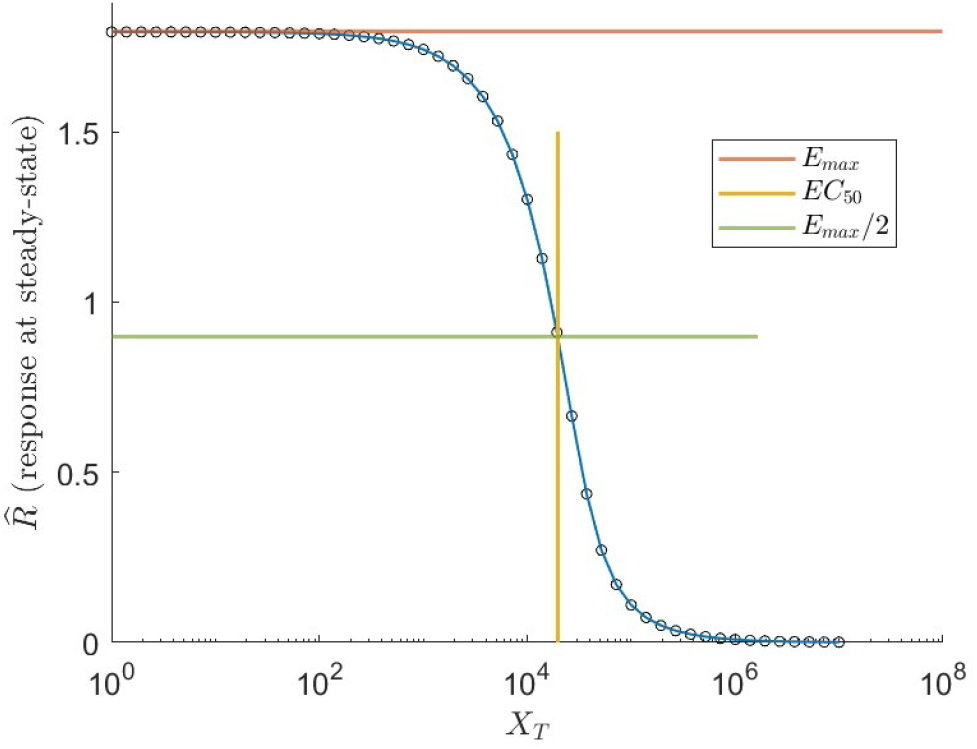
*KPC* model. The response at steady-state, 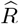, as a function of total ligand (solid blue line) was obtained using the general equation for 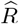 (Eqn. (86) in the Appendix). Black circles correspond to values of *R*(*t*) obtained with the ODE system at *t* = 2000 *s* and a wide range of values of *L*_*T*_. The horizontal red line shows the value of *E*_max_ computed with Eqn. (89) while the green line corresponds to *E*_max_*/*2. The vertical yellow line corresponds to *EC*_50_ obtained with equation Eqn. (90). Parameter values: *k*_*on*_ = 10^*−*5^ *s*^*−*1^[]^*−*1^, *λ* = 0.04 *s*^*−*1^, *k*_off_ = 0.05 *s*^*−*1^, *k*_*p*_ = 0.09 *s*^*−*1^, *k*_+_ = 0.1 *s*^*−*1^, *k*_*−*_ = 0.05 *s*^*−*1^, *T*_*T*_ = 3 *×* 10^4^ TCRs/cell. The symbol [] indicates a concentration unit (number of molecules per cell) for membrane and cytoplasmic molecules.

### 3.13 Modified ST model (*ST-mod*)

The serial triggering concept (*ST*) was originally proposed by Valitutti and Lanzavecchia in [26], and an extended mathematical model was proposed five years later [13] defined by the following ODE system:

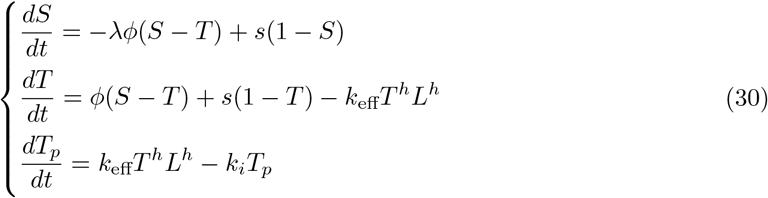

According to the description of this model [13], two subsets of TCRs are distinguished: one within a so-called interface pool and the other within a spare pool. TCRs within the interface pool (denoted *T*) are situated in an area, determined by the membrane interface between a T cell and an APC, where they can interact with pMHC molecules. These interactions, characterised by an effective kinetic rate *k*_eff_ and a kinetic order *h >* 1, result in the formation of triggered (phosphorylated) TCRs (denoted *T*_*p*_), which are subsequently internalised with constant rate *k*_*i*_ [13]. Conversely, the spare pool (denoted *S*) contains TCRs located outside the interface area, with the ratio of interface area to spare area characterized by the parameter *λ*. Free TCRs diffuse between the two pools with constant rate *ϕ* per unit membrane area, and undergo turnover between the membrane and the cytoplasm with constant rate *s* per unit membrane area. At close inspection, however, it can be seen that Eqns. (30) do not correctly translate into mathematical expressions the conceptual description of the model. For instance, according to those equations, in the absence of ligand each pool ends up accumulating half of the total TCRs, independently of the ratio *λ* and of the initial values of *S* and *T*.

Here we propose a modification of the above equations, in an attempt to make them correspond accurately to the mechanism conceptually described in [13]. In this model variant, the rate of TCRs turnover between the membrane and the cytoplasm is denoted *σ*, and all other parameters are denoted as in the original model. This reformulation introduces a new definition for the flow between areas, represented by the first term in the equations for *dS/dt* and *dT/dt*. Additionally, it redefines the input rate of receptors, which is now proportional to the respective areas of the two pools. The new equations, with the state-variables adimensionalyzed with respect to the total amount of TCRs in the absence of ligand, that is, with respect to *S*_0_ + *T*_0_, are the following:

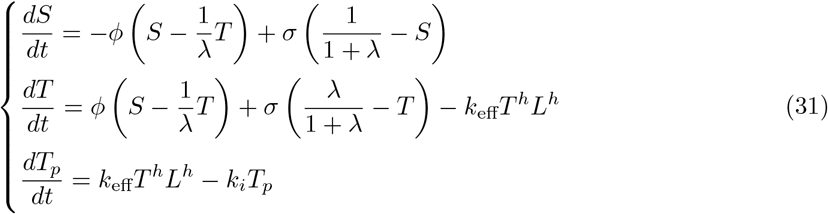

In this model the response is defined by *R*(*t*) = *T*_*p*_(*t*). Notice that, according to the definition of *λ*, it verifies *λ* = *T*_0_*/S*_0_. We refer to this variant model as modified ST or *ST-mod*. Note, however, that both models, *ST* and *ST-mod*, share the same graphical scheme (see Fig. 1, panel 12).

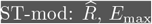 and *EC*_50_.

For *h* = 1:

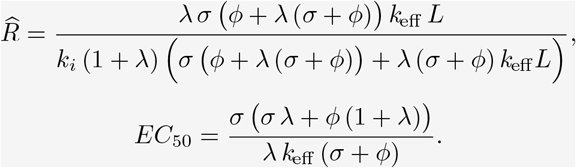

For *h* = 2:

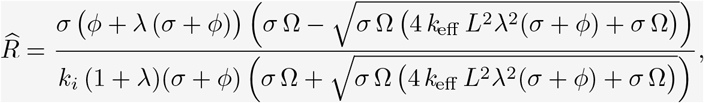

with:

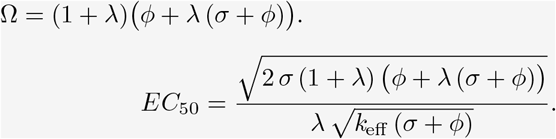

For *h ≥* 1 (*h ∈*ℕ):

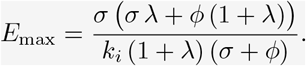

Due to the complexity of the calculations we were unable to obtain a general expression of *EC*_50_ for *h* = 3, 4.

## 4 Conclusions

In this work we have contributed to filling a gap in the literature, by deriving the expressions of maximum drug effect (*E*_max_) and half-maximal effective concentration (*EC*_50_) for the main T-cell activation models for which those expressions were previously unknown. Knowledge of these expressions is necessary to perform analyses of key model properties, including identifiability, observability, and sensitivity. Thus, the results presented in this paper enable a deeper analysis of the existing models of T-cell activation, and facilitate obtaining biological insights about the involved mechanisms.

Our results have revealed a clear trend. While the previously known expressions of *E*_max_ and *EC*_50_ were relatively simple, the formulas of these quantities become more involved as the complexity of the models increases. As a result, in some models the impact of specific parameters on the maximum drug effect and half-maximal effective concentration is less direct, and possibly easier to be compensated by changes in other parameters. This, in turn, implies that inferring the values of such parameters from measurements of *E*_max_ and *EC*_50_ is likely to be quite difficult, if not impossible. Since those are often the only measurable quantities, this fact hints at the possible existence of identifiability issues in the corresponding models. We are currently investigating this aspect, and the results will be presented elsewhere.

## Appendix: derivations of 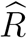, *E*_max_ and *EC*_50_ formulas

### KPR sustained signalling (KPR-SS)

*Proof*. **Step 1:** The ODE system at steady-state:

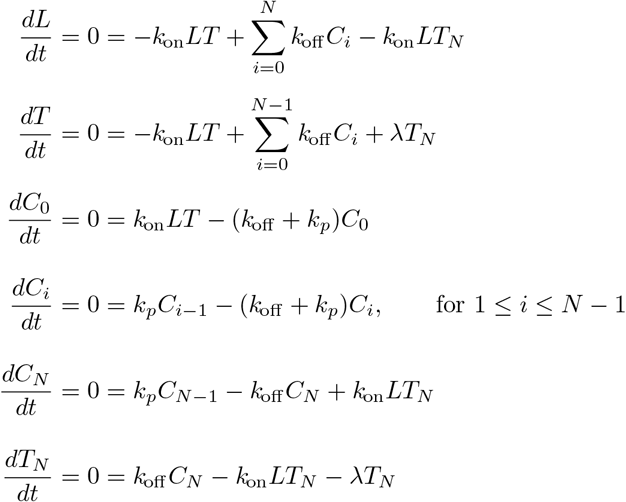

Conservation equations, with 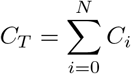:

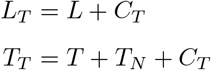

**Step 2:** In this model the response is defined as 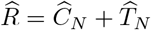. Firstly, we should express 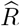 in terms of only one variable, 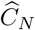 or 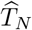. Thus, from the equation for *dT*_*N*_ */dt* at steady-state and using the conservation equation for 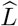,

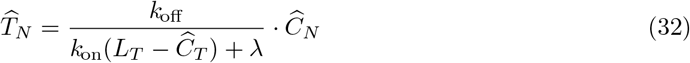

In order to find an expression for 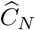 in terms of 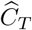 we proceed as follows. First, from Eqn. *dC*_*i*_*/dt* one has,

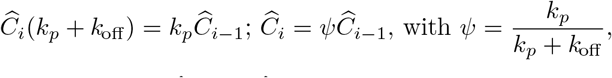

and applying it recursively it yields 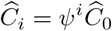, for *i* = 0, …, *N −* 1. Therefore:

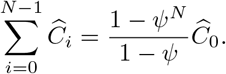

On the other hand,

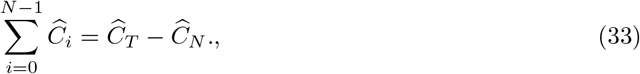

hence:

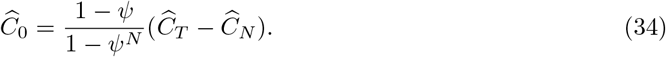

Summing the steady-state equations for *dT/dt* and *dC*_0_*/dt*, and using Eqns. (32) and (34) yields,

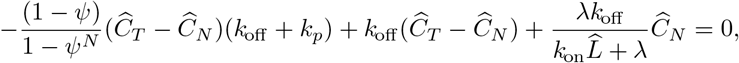

and taking into account that 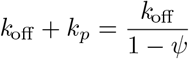 one has,

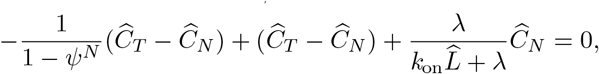

which, after some algebraic rearrangements, yields:

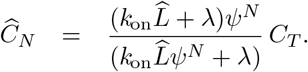

And inserting there the conservation equation for 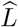, yields:

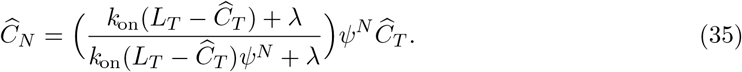

Now, inserting Eqns. (32) and (35) in the definition of *R* one has,

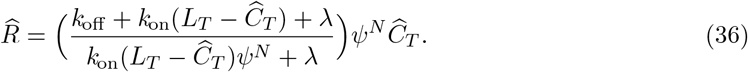

In order to obtain an expression for 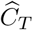 in terms of *L*_*T*_ and *T*_*T*_, it suffices to follow the procedure outlined in subsection 2.3.2 and substitute the conservation equations into the steady-state equation for *dL/dt*:

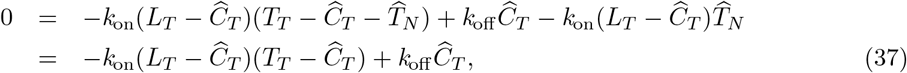

hence,

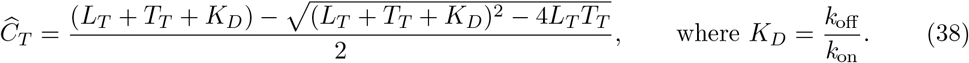

Finally, substituting Eqn. (38) into Eqn. (36) gives a closed-form expression for *R* in terms of *L*_*T*_ and *T*_*T*_.

**Step 3:** To calculate 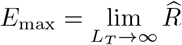 we make use of Eqn. (37) and proceed as in subsection 2.3.2:

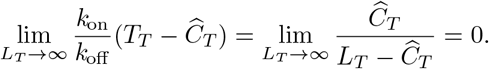

Consequently, 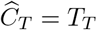 in that limit; and applying this limit also to Eqn. (36) one has:

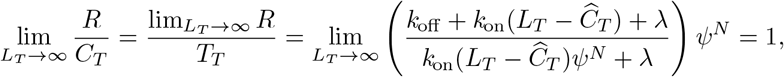

therefore:

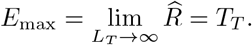

**Step 4:** In this model, *EC*_50_ is the value of *L*_*T*_ for which it verifies 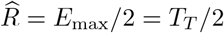, that is, it verifies:

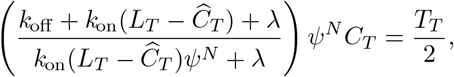

with *C*_*T*_ given by Eqn. (38).

Solving this equation for *L*_*T*_ gives the following closed-form expression for *EC*_50_ that was previously unavailable in the literature:

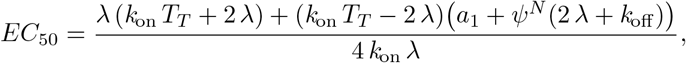

with,

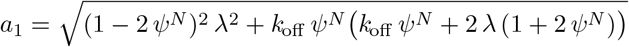

□

### KPR with induced rebinding (KPR-IR)

*Proof*. **Step 1:** The ODE system at steady-state:

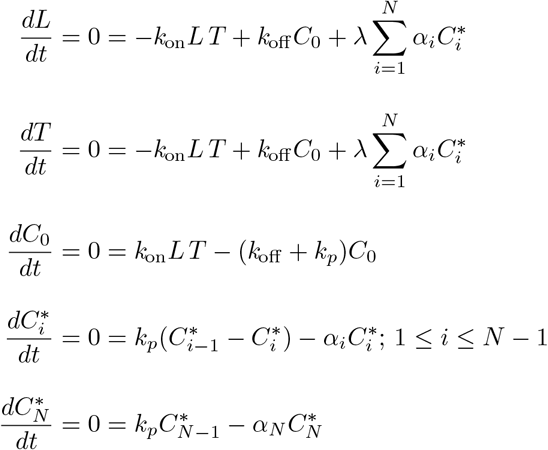

Conservation equations, with 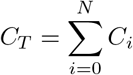:

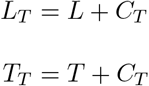

**Step 2:** In this model the response is given by 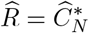. From the steady-state equations for 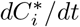 and 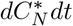 we get,

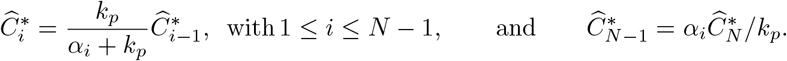

This leads to,

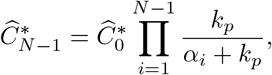

and

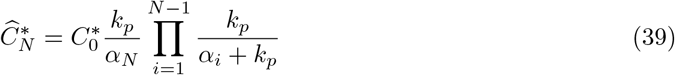

This allows to obtain the following expression for 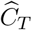:

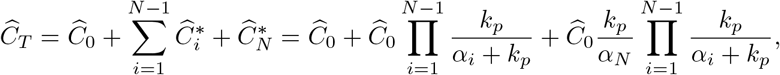

and, therefore,

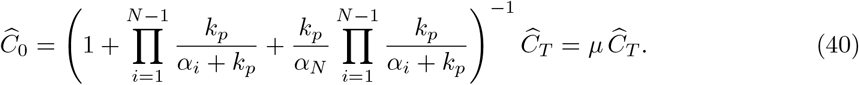

with 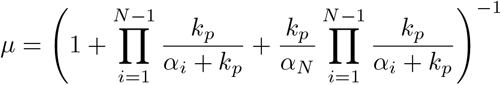.

Introducing this last result into Eqn. (39) yields,

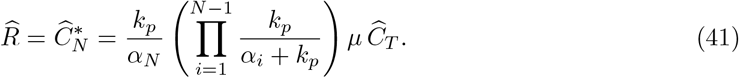

To express 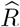 in terms of only parameters and constants we need to express 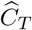 in terms of *L*_*T*_ and *T*_*T*_. For that, it suffices to introduce Eqn. (40) into the steady-state equation for *dC*_0_*/dt* giving,

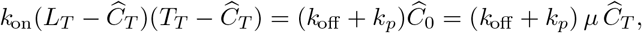

and solving that equation for 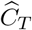 yields,

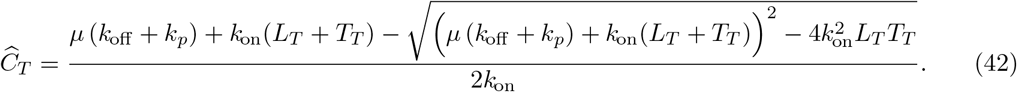

**Step 3:** From first principles we have 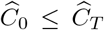. Now, from the equation for *dC*_0_*/dt* at steady-state one has:

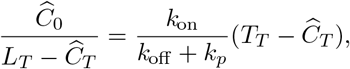

and in the limit *L*_*T*_ *→ ∞* it verifies:

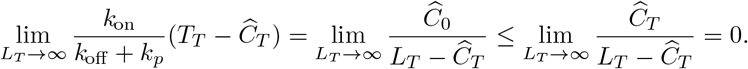

Consequently, in that limit 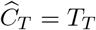. Inserting this result into Eqn. (41), we find that:

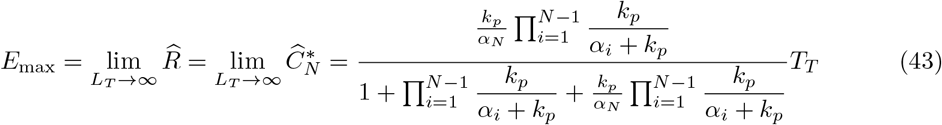

**Step 4:** *EC*_50_ can be determined by finding the concentration of *L*_*T*_ that satisfies the expression 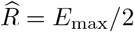. Taking into account Eqns. (41) and (43), this implies finding the concentration of *L*_*T*_ that satisfies:

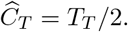

Substituting there *C*_*T*_ by its value defined in Eqn. (42), and after some algebraic rearrangements, it yields:

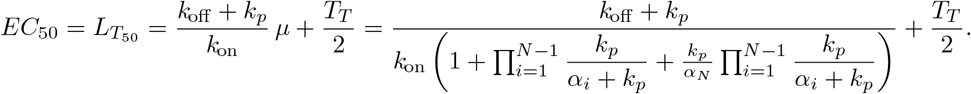

□

### KPR with limited signalling and incoherent feed-forward loop (*KPR-LS-IFF*)

*Proof*. **Step 1:** The ODE system at steady-state, with 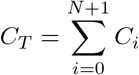, is:

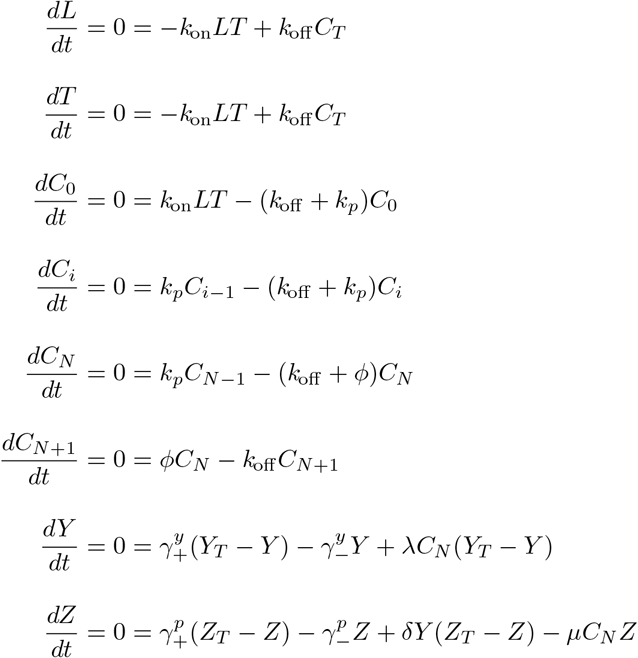

Conservation equations:

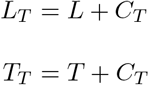

**Step 2:** The response in this model is defined by *R*(*t*) = *P* (*t*). To establish a relationship between the total number of complexes and the variable *P* (*t*), we first obtain an expression for the total number of complexes at steady-state. For that we proceed as in subsection 2.3.2, and using the steady-state equation for *dL/dt* and the conservation equations, we get:

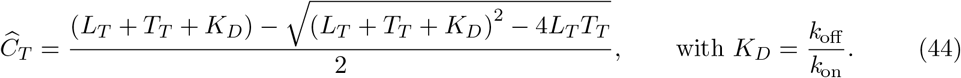

Next, we seek to express 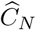 in terms of 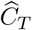. For that we first notice that by definition:

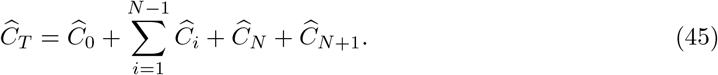

Following again the procedure in subsection 2.3.2 and using the steady-state equations for *dC*_*i*_*/dt* we have 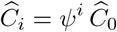, where 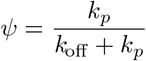. Therefore,

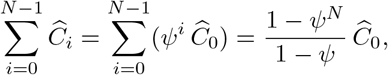

Also, from the equation for *dC*_*N*+1_*/dt* we have:

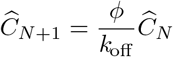

Last, adding the steady-state equations for *dC*_0_*/dt* and *dL/dt*, leads to:

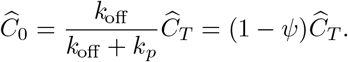

Replacing the above three results in Eqn. (45) yields:

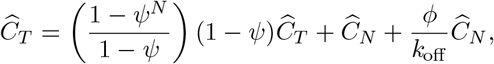

and solving for 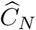 gives:

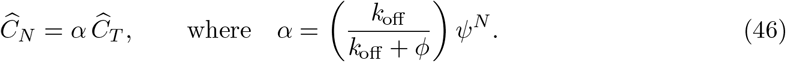

Now, from the equations for *dY/dt* and *dZ/dt* at steady-state we have:

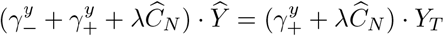

and

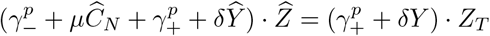

From the first of these two equations we have:

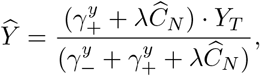

and inserting this result into the second equation we get:

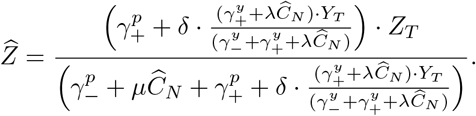

Finally, using Eqn. (46), we have:

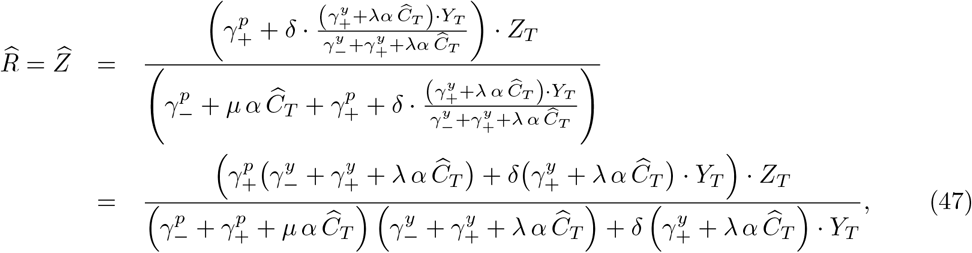

with:

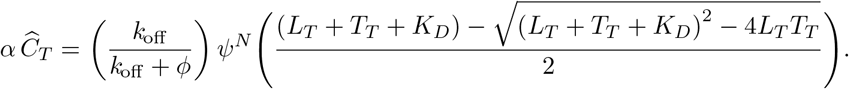

**Step 3:** In this model, the response 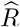 as a function of *L*_*T*_ displays, for many reasonable parameter values, a bell-shaped behaviour (see Fig. 3 in the main text), therefore, in general *E*_max_ does not correspond to 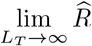.

To obtain *E*_max_ one has to proceed in two steps, first calculate 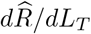 using Eqn. (47), and then calculate *L*_*T*_ from the equation 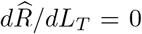. Denoting 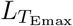 this particular value of *L*_*T*_, one has:

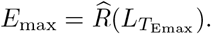

According to that procedure, we obtain the following equation for 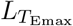:

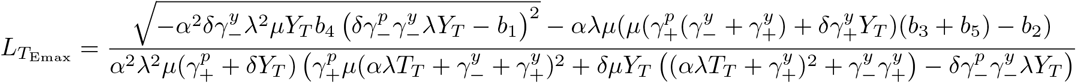

where:

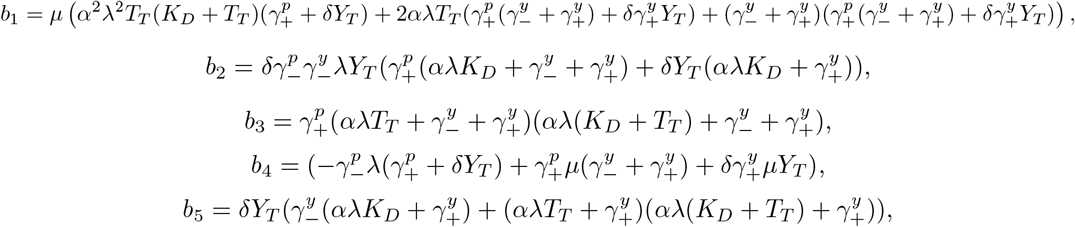

with *α* defined in Eqn. (46).

Substituting this result into Equation (47) yields an expression for *E*_max_. Owing to its big size, however, this expression is only available in our online repository (https://github.com/Xabo-RB/Inmunology-analysis.git).

**Step 4:** *EC*_50_ is the ligand concentration at which 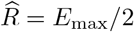, that is, it is the value of *L*_*T*_ that verifies:

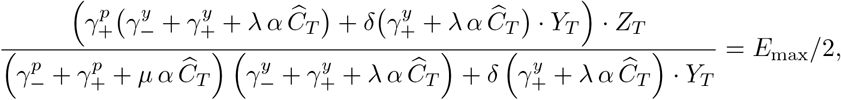

with 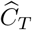 given by Eqn. (44). As for the case of *E*_max_, due to its big size, the complete expression for *EC*_50_ is available only in our online repository. □

### KPR with negative feedback (*KPR-NF1*)

*Proof*. **Step 1:** The ODE system at steady-state, with 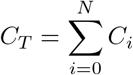, is:

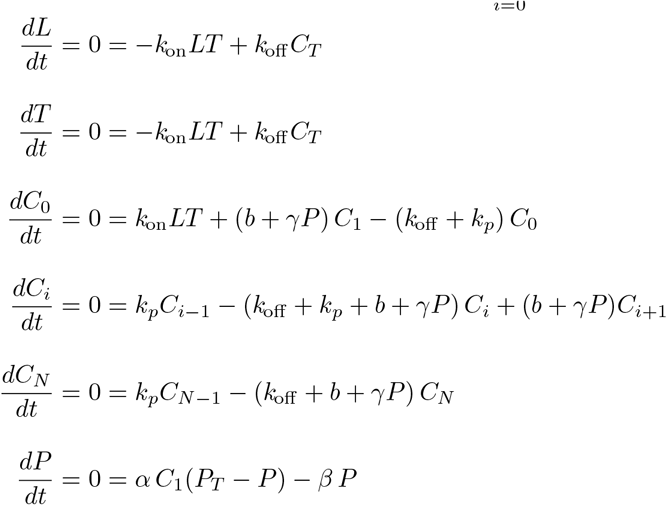

The conservation equations are:

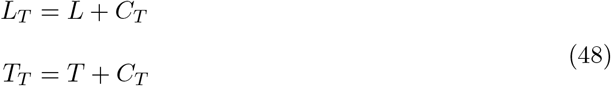

**Step 2:** We have to relate the response of the system, defined in this model as *R*(*t*) = *C*_*N*_ (*t*), to the total number of complexes, *C*_*T*_. For that we need to proceed in a way different from the previous models. First, from the steady-state equations for *dC*_*i*_*/dt*, we have:

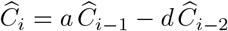

and

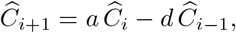

with 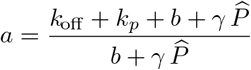 and 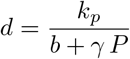.

Writing those equations in vector notation for *i* = *N −* 1:

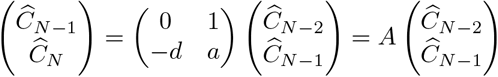

Proceeding recursively, it yields:

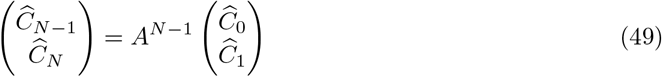

On the other hand, matrix *A* can be diagonalised to give:

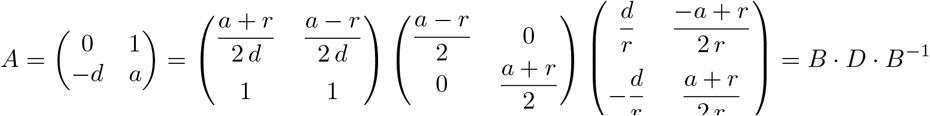

where 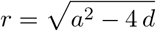. Therefore:

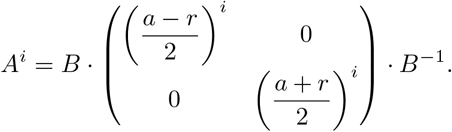

From Eqn. 49, we get:

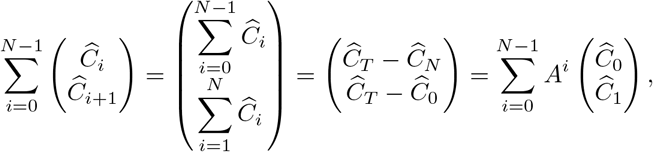

and using the following known expression for the geometric sum of a diagonalizable matrix:

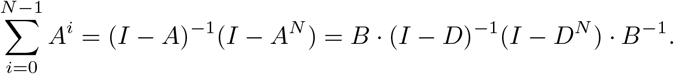

Then, after some straightforward matrix operations, we have:

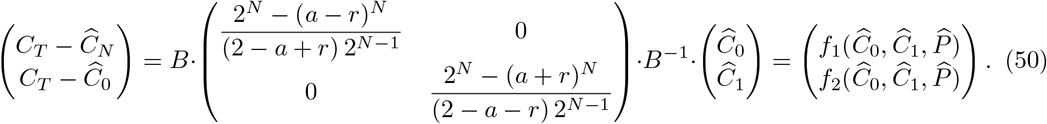

Therefore, from the above equation we have:

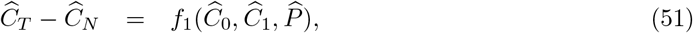

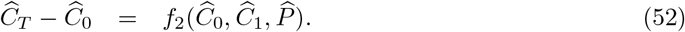

On the other hand, from the steady-state equations for *dL/dt* and *dT/dt* the following expressions for 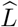 and 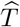 in terms of *L*_*T*_ and *T*_*T*_ can be obtained:

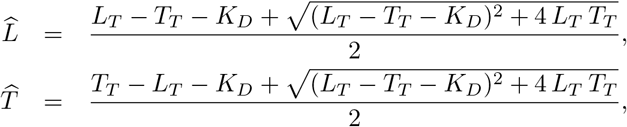

with *K*_*D*_ = *k*_on_*/k*_off_; and proceeding as in subsection 2.3.2 from the steady-state equation for *dL/dt* one has:

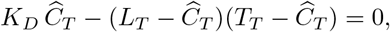

and therefore:

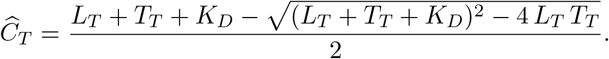

We need now to express 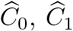 and 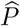 in terms of 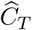, which will link them directly through the above equation to *L*_*T*_ and *T*_*T*_. For that we first use the steady-state equation for *dP/dt* to get:

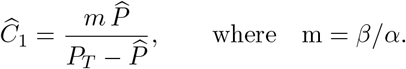

Next, we turn to the steady-state equation for *dC*_0_*/dt*:

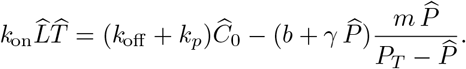

Solving this equation for 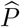 yields:

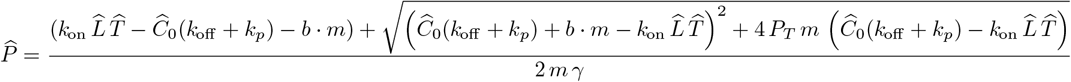

After having obtained an expression for 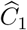 and 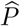 in terms of 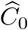, we go on to obtain an expression for 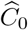 in terms of 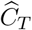. For that we notice that Eqn. (52) can be written as:

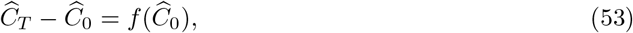

where 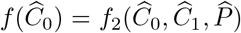 with 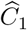 and 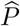 given by the above equations. Solving Eqn. (53) for 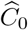 and inserting the obtained expressions for 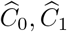 and 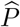 into 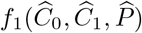 in Eqn. (51) leads to:

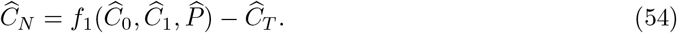

Hence, since in this model 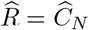, we obtained the expression for 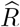 we were looking for.

A drawback of this result is that as *N* increases the formulas become exceedingly large, so that even for *N* = 3 one has to resort to numerical calculations.

**Step 3:** In references [3, 4, 18]it was mentioned that in this model the response 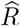 as a function of *L*_*T*_ exhibits a bell-shaped behaviour. We have confirmed by numerical calculations that this is the case for many parameter combinations and more than two kinetic steps (*N >* 2). Therefore, like with the KPR-LS-IFF loop model, in this model *E*_max_ does not correspond in general to 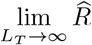. Instead, we have to specify a particular set of parameter values, leaving *L*_*T*_ as a free parameter, and define numerically 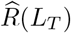. This allows to obtain the peak of this function for each particular set of parameters.

**Step 4:** *EC*_50_ can be determined by finding the concentration of *L*_*T*_ that satisfies the expression: 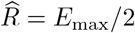. Because in this model a general expression for *E*_max_ cannot be obtained, the value of *EC*_50_ has to be calculated numerically, which requires defining a particular set of parameter values. □

### KPR with negative feedback II (*KPR-NF2*)

*Proof*. Given that the structure of this model is identical to that of the *KPR-NF1* model, all results from Eqn. (48) to (54) hold also in this model. However, in this model the response is defined as:

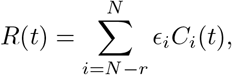

with *ϵ*_*i*_ *< ϵ*_*i*+1_, for *i* = *N − n* to *N −* 1, and *ϵ*_*N*_ = 1.

The procedure developed for the KPR-NF1 model allows to calculate 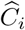 from *i* = 1 to *N* and, therefore, an expression for 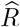 can be obtained also in this model (albeit a quite complex one) once *n* and *α*_*i*_ are specified. □

### KPR stabilizing activation chain (*KPR-SAC*)

*Proof*. **Step 1:** The ODE system at steady-state is:

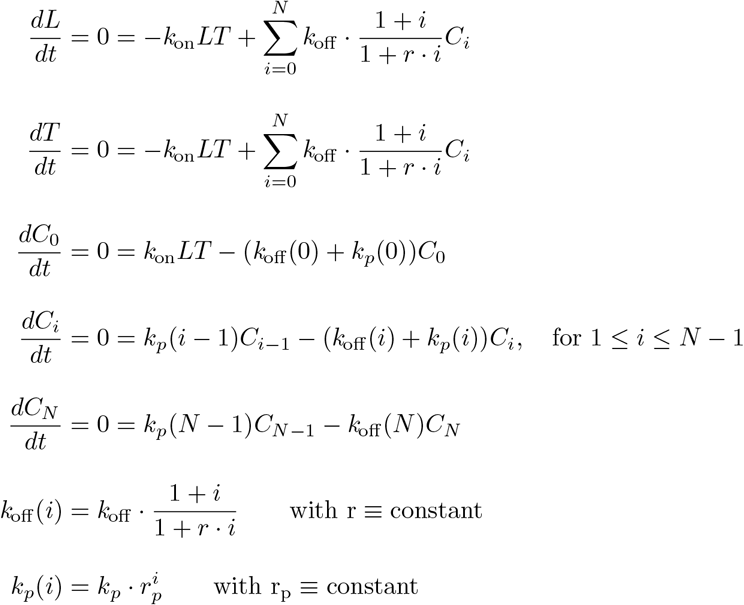

Conservation equations, with 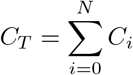:

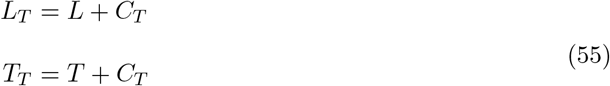

**Step 2:** In this model the response is defined as *R*(*t*) = *C*_*N*_ (*t*). In order to calculate *C*_*N*_ at steady-state, that is, 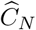, we proceed as follows. We first substitute *k*_*p*_(*i*) and *k*_off_(*i*) with their corresponding expressions in the equations for *dC*_*i*_*/dt* and *dC*_*N*_ */dt* at steady-state:

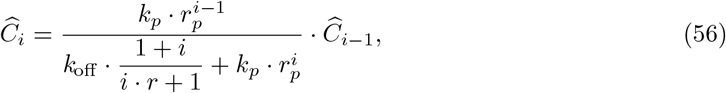

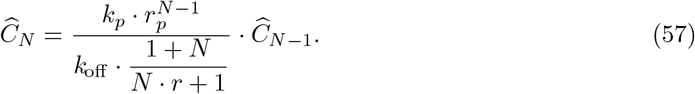

To simplify the notation we define:

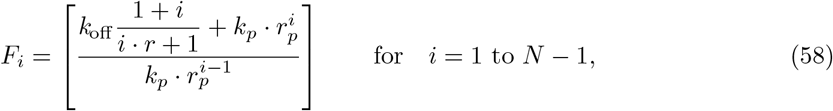

and

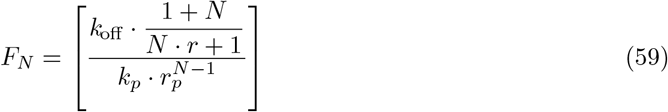

Applying induction to Eqn. (56) one gets:

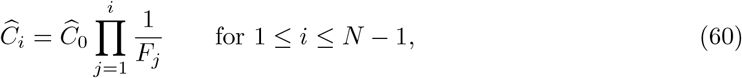

and, in particular:

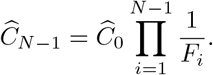

Inserting the above equation into Eqn. (57) yields:

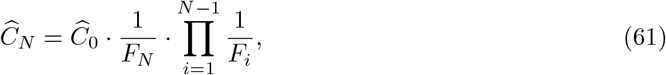

and isolating variable 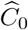 we get,

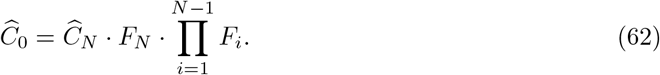

The total number of complexes is given by: 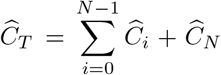. Therefore, inserting here Eqns. (60) and (61), and after some algebraic rearrangement, we find:

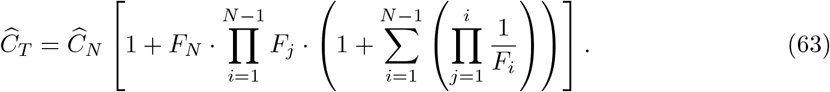

Using now the straightforward result:

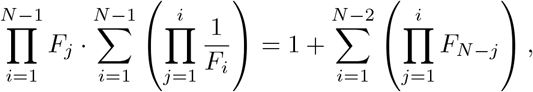

we can rewrite Eqn. (63) in this way:

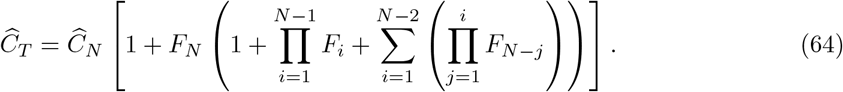

Finally, since in this model *R*(*t*) = *C*_*N*_ (*t*), we have at steady-state:

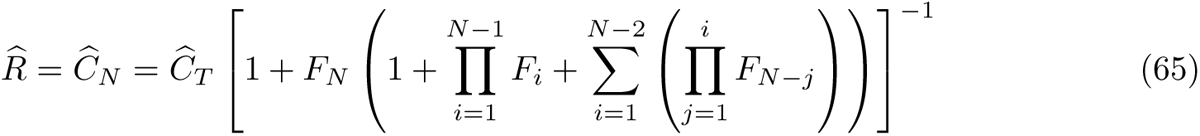

An expression for *C*_*T*_ can be derived using the steady-state equation for *dC*_0_*/dt* and inserting there Eqns. (62) and (65). In this way it is obtained:

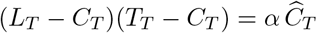

Therefore,

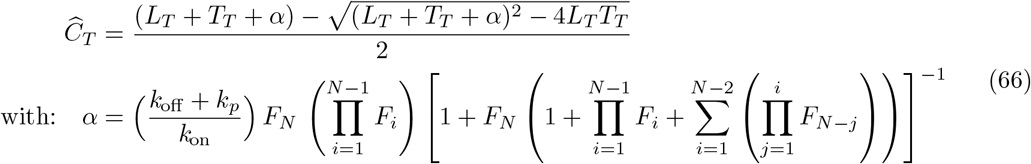

**Step 3:** Similarly to previous models, from the equation for *dC*_0_*/dt* at steady-state and the conservation equations, and considering that 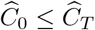, one has:

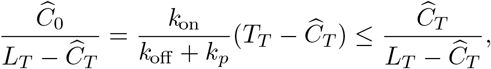

thus:

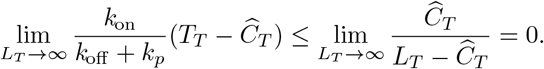

Therefore, in that limit it verifies 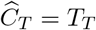. Hence:

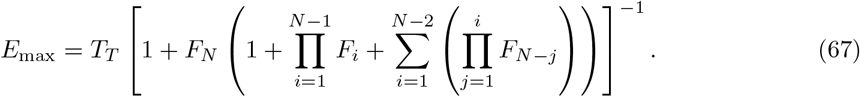

**Step 4:** *EC*_50_ can be determined by finding the concentration of *L*_*T*_ that satisfies the expression *R* = *E*_max_*/*2. From equations (65) and (67), we obtain that this condition is satisfied for:

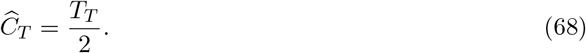

Inserting Eqn. (61) into Eqn. (64) yields:

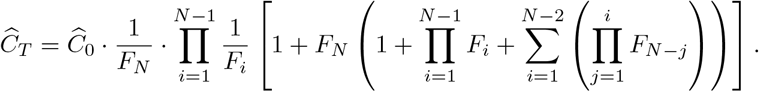

Let us simplify notation again by defining:

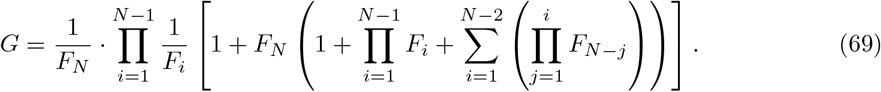

Then,

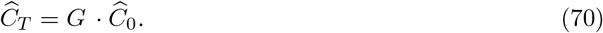

From the equation for *dC*_0_ = */dt* at steady-state, and using the above equation, we have:

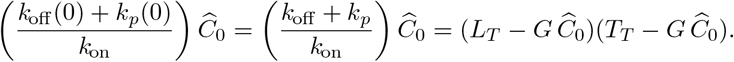

Hence,

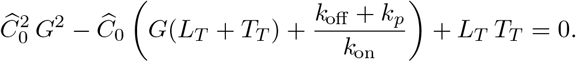

Therefore,

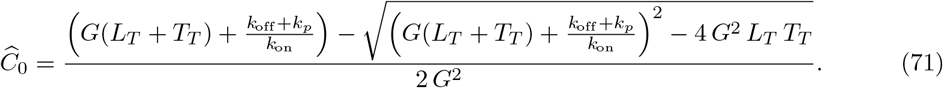

Inserting Eqns. (70) and (71) into Eqn. (68) yields:

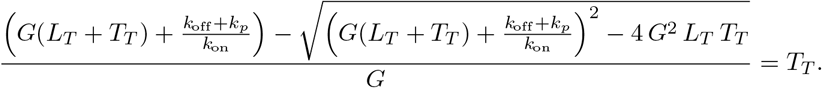

Isolating now *L*_*T*_ from the above equation gives:

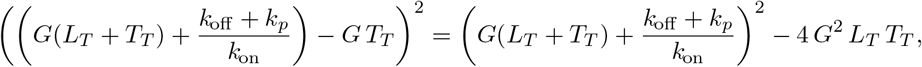

and, after some algebraic rearrangements, we get:

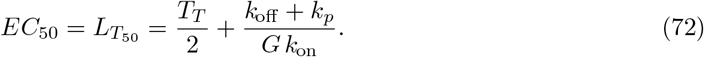

□

### Zero-order ultrasensitivity model (*ZU*)

*Proof*. **Step 1:** The ODE system at steady-state is:

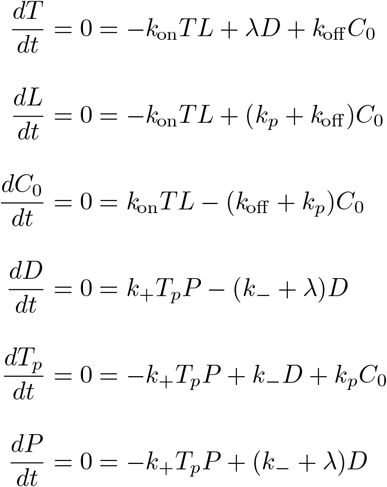

The conservation equations are:

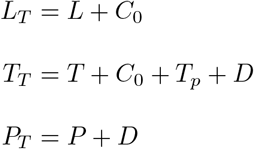

**Step 2:** In this model the response is defined as *R*(*t*) = *T*_*p*_(*t*). Thus, obtaining an expression for *R* is equivalent to find an expression for *T*_*p*_. For that we first notice that from the conservation equation for *L* we have:

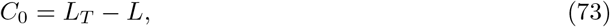

and from the equations for *dT/dt* and *dC*_0_*/dt* at steady-state we have, respectively:

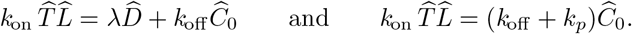

Combining them and using Eqn. (73) yields:

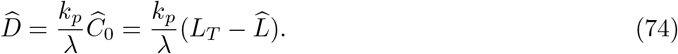

On the other hand, from Eqn. (74) and the conservation equation for *P* it follows,

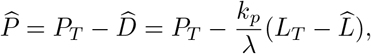

Finally, using this expression in the equation for *dP/dt* at steady-state, we are lead to:

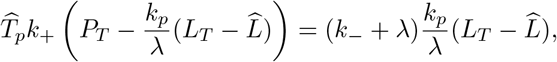

and hence,

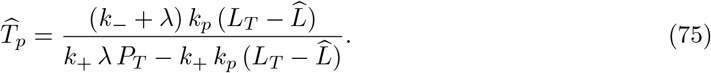

**Step 3:** To calculate 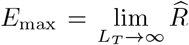 we note that *C*_0_(*t*) *≤ T*_*T*_ and, therefore, 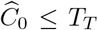. Now, from the equation for *dL/dt* at steady-state, and using the conservation equations, we obtain:

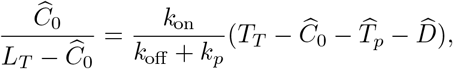

and in the limit *L*_*T*_ *→ ∞* it verifies:

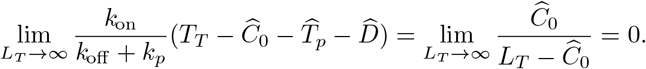

Consequently, 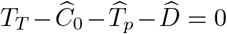, and using the first result from Eqn. (74), one has in that limit: 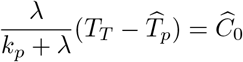. Combining with the conservation equation, 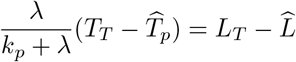.

Using this result within Eqn. (75) yields,

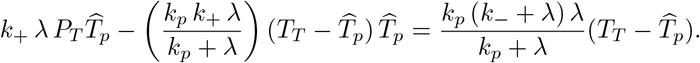

Rearranging terms, this expression can be written as:

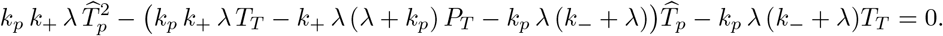

Finally, denoting

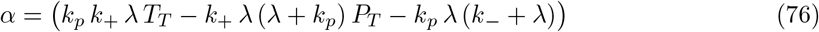

it is obtained:

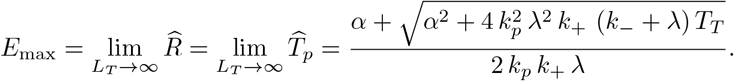

**Step 4:** To derive an expression for *EC*_50_ it is necessary to first express 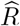, and hence 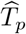, in terms of *L*_*T*_. For that, we start from the equation for *dL/dt* at steady-state together with the conservation equations and Eqns. (74) and (75):

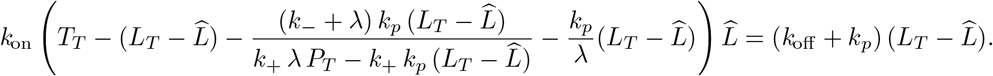

This gives the following cubic polynomial in 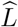:

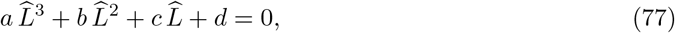

where

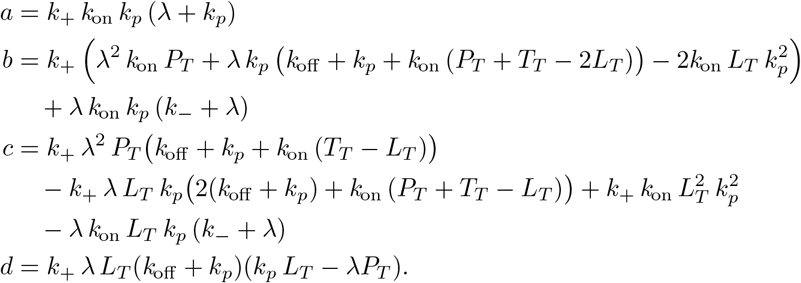

Of the three solutions to this polynomial we are interested in the one that fulfils the following two conditions: 1) it is positive real, and 2) it is lower than *L*_*T*_. Inserting this solution into Eqn. (75) gives 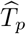, and hence 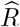, in terms of *L*_*T*_.

Finally, isolating *L*_*T*_ from the equation 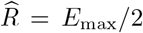 gives 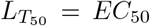. Nevertheless, we cannot show here a general formula because the expressions for both 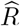 and *EC*_50_ are exceedingly long. Instead, the complete expression for *EC*_50_ is available in our online repository. □

### KPR with zero-order ultraspecificity (*KPZU*)

*Proof*. **Step 1:** The ODE system at steady-state, with 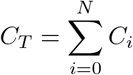, is:

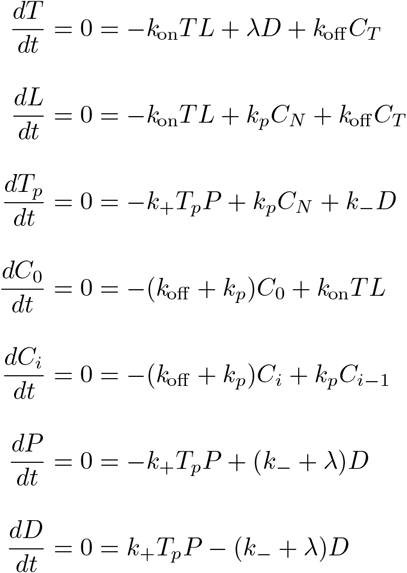

The conservation equations are:

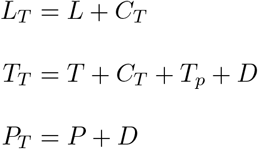

**Step 2:** Like in th ZU model, in this model the response is also defined as *R*(*t*) = *T*_*p*_(*t*). Thus, in order to obtain an expression for *R* it is enough to find an expression for *T*_*p*_. This model includes a simple kinetic proofreading chain in addition to the processes of the ZU model, such that in the steady-state there is the following simple relationship between complexes *C*_*i*_ and *C*_*i−*1_:

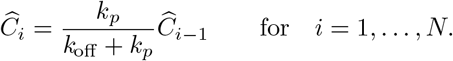

Thus, applying it recursively, one has:

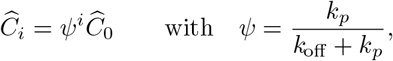

and, in particular:

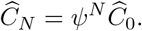

From here it is obtained:

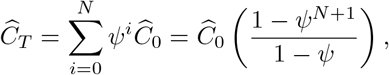

and, therefore:

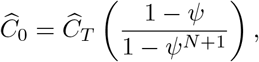

and

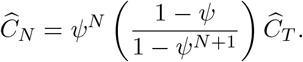

Summing the equations for *dT/dt* and *dD/dt* at steady-state yields:

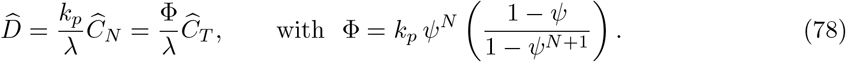

and using this result in the conservation equation for 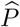 one has:

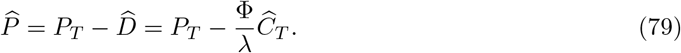

On the other hand, from the equation for *dD/dt* at steady-state one has:

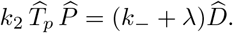

Substituting here 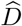 and 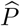 as given by Eqns. (78) and (79) yields:

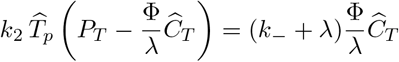

Therefore,

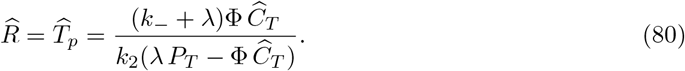

Notice that in Eqns. (78)–(80) and in the following calculations the complex parameter Φ plays the same role as *k*_*p*_ in the ZU model.

**Step 3:** To calculate 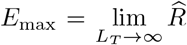 we note that *C*_0_(*t*) *≤ C*_*T*_ and, therefore, 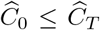. Now, from the equation for *dC*_0_*/dt* at steady-state, and using the conservation equations, we obtain:

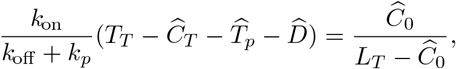

and in the limit *L*_*T*_ *→ ∞* it verifies:

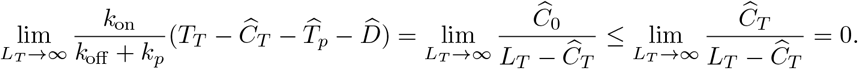

Consequently, in that limit 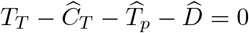, and using Eqn. (78), one has:

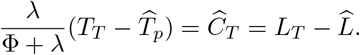

Using this result within Eqn. (80) yields,

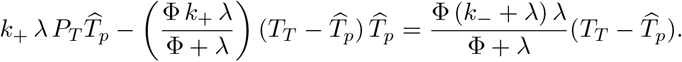

Rearranging terms, this expression can be written as:

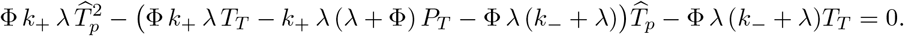

Finally, denoting *α* = (Φ *k*_+_ *λ T*_*T*_ *− k*_+_ *λ* (*λ* + Φ) *P*_*T*_ *−* Φ *λ* (*k*_*−*_ + *λ*)) it is obtained:

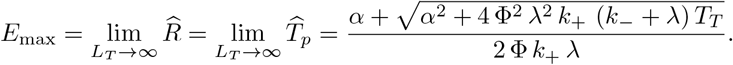

**Step 4:** To derive an expression for *EC*_50_ it is necessary to first express 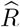, and hence 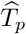, in terms of *L*_*T*_. For that, we start from the equation for *dT/dt* at steady-state together with the conservation equations and Eqns. (78) and (80):

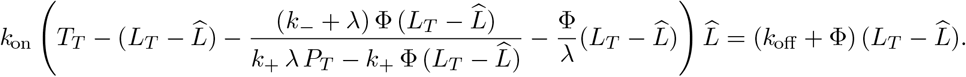

This gives the following cubic polynomial in 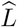:

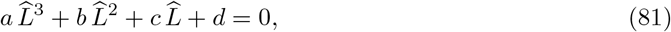

where:

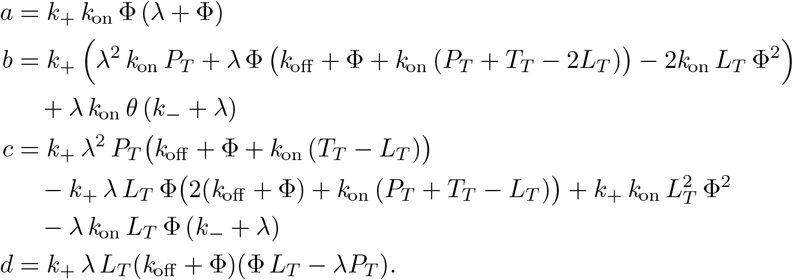

Of the three solutions to this polynomial we are interested in the one that fulfills the following two conditions: 1) it is positive real, and 2) it is lower than *L*_*T*_. Then, substituting 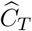 for 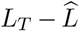 in Eqn. (80) and inserting there the above solution for 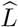 gives 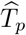, and hence 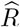, in terms of *L*_*T*_.

Finally, isolating *L*_*T*_ from the equation 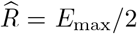 gives 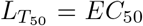. Nevertheless, like in the ZU model, we cannot provide here a general formula because the results for both 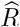 and *EC*_50_ involve quite big expressions. Therefore, in order to obtain values for these quantities, one has to resort to numerical calculations using specific sets of parameter values. □

### KPR with concentration compensation (*KPC*)

*Proof*. **Step 1:** The ODE system at steady-state, with 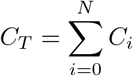, is:

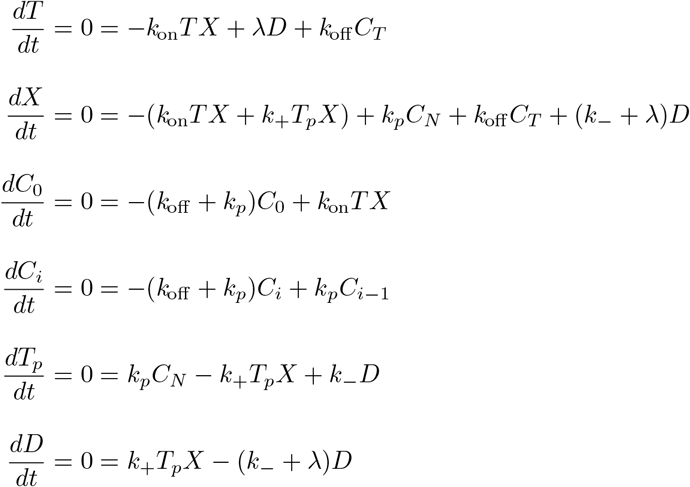

The conservation equations are:

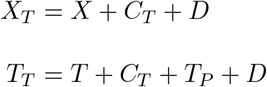

**Step 2:** Similarly to the ZU and KPZU models, in this model the response is also defined as *R*(*t*) = *T*_*p*_(*t*). We are, therefore, interested in finding an expression for *T*_*p*_ in terms of *L*_*T*_. For that we will proceed in a way similar to the KPZU model, noticing that this model also includes a kinetic proofreading chain. Thus, from the equation for *dC*_*i*_*/dt* at steady-state one has:

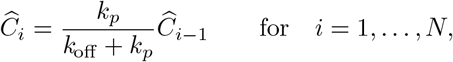

and applying it recursively, it yields:

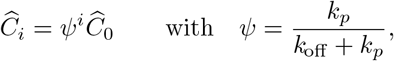

and:

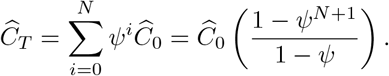

Therefore:

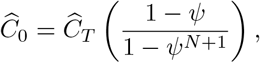

and

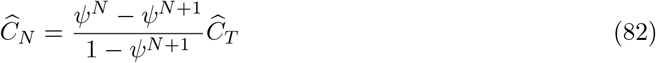

Summing the steady-state equations for *dT/dt* and *dD/dt* and using the above equation for 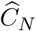 yields:

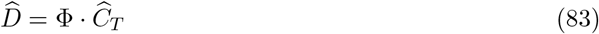

where 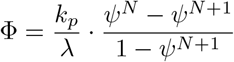.

Inserting the conservation equations and Eqn. (83) into the equations for *dT/dt* and *dD/dt* at steady-state yields, respectively,

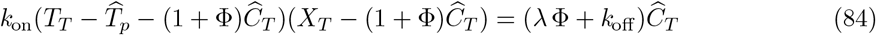

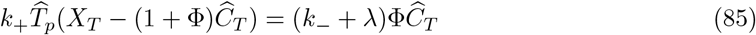

Finally, solving the system formed by Eqns. (84) and (85) for 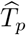 and 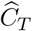, we get:

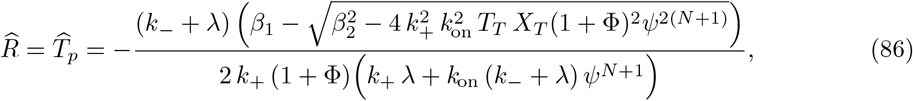

where:

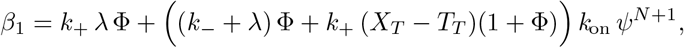

and

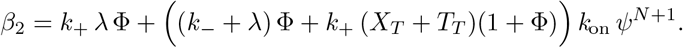

**Step 3:** To calculate 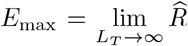 we note that, as in previous models, *C*_0_(*t*) *≤ C*_*T*_ (*t*) and, therefore, 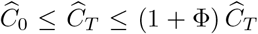. Now, from the equation for *dC*_0_*/dt* at steady-state, and using the conservation equations, we obtain:

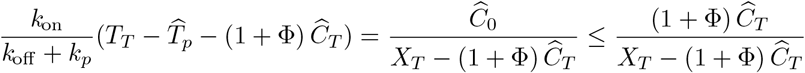

In the limit when there is ligand saturation,

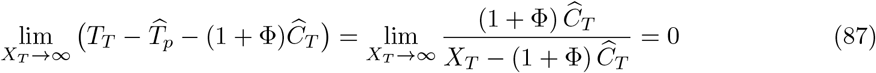

On the other hand, from the equation for *dD/dt* at steady-state, and using the conservation equations and Eq. (83) one has,

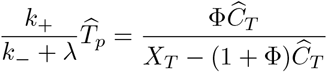

Taking now the limit given when there is ligand saturation, we find:

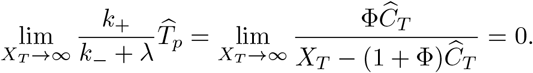

Therefore, 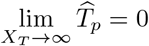. Notice that this result is at odds compared to most of the other models.

Introducing this last result into Eqn. (87) we get,

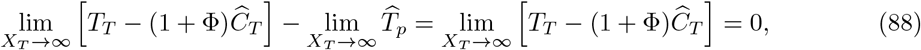

therefore,

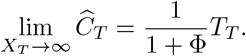

This conclusion is validated numerically as shown in Fig. 5 in the main text. In addition, Eqn. (88) indicates that, under conditions of ligand saturation, 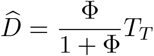.

Analysing Eqn. (86) it can be shown that 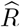 increases as *L*_*T*_ decreases. This indicates that *E*_max_ is obtained in the limit 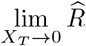. Therefore, taking *X*_*T*_ = 0 in Eqn. (86) (that is, in the auxiliary parameters *β*_1_ and *β*_2_) one has:

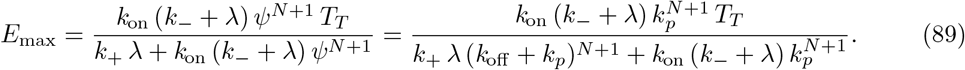

**Step 4:** *EC*_50_ is the ligand concentration that satisfies the relation: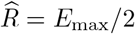, with 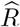 given by Eqn. (86) and *E*_max_ by Eqn. (89). Thus, according to those equations, *EC*_50_ is the ligand concentration that satisfies:

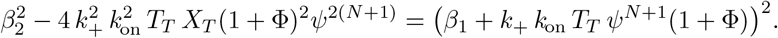

Then, after after substituting there the auxiliary parameters *β*_1_ and *β*_2_, and solving the resulting equation for *X*_*T*_ and simplifying it, it is obtained:

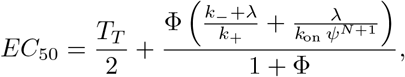

which, after substituting *ψ* and Φ, and after some algebraic rearrangements, yields:

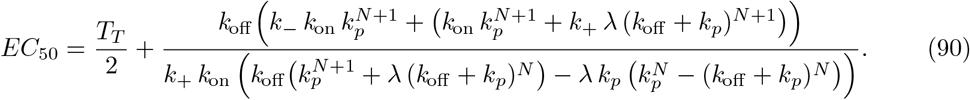

□

### Modified serial triggering (ST-mod)

*Proof*. **Step 1:** The ODE system at steady-state is:

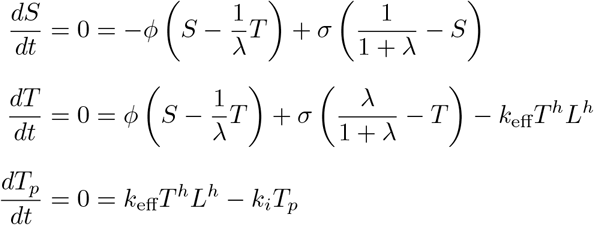

This model is an extension of a previous one proposed by Bachmann *and col* [27] and inherits from it the following assumptions: (1) ligand-receptor complexes and complex multimers are in quasi steady-state relative to the kinetics of TCRs, and (2) the concentration of ligand-receptor complexes is much smaller than that of TCRs. Therefore, in this model the ligand concentration or density is considered a parameter, denoted here *L*. The three state variables of this model, *S, T* and *T*_*p*_, denote the density of TCRs in three different compartments in a T lymphocyte membrane, respectively, TCRs in the spare pool, free TCRs in the interphase pool, and triggered or activated TCRs, which are marked for internalization and degradation.

**Step 2:** The response is quantified here in terms of activated TCRs, that is, 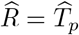.

From the steady-state equation for *dA/dt* one has:

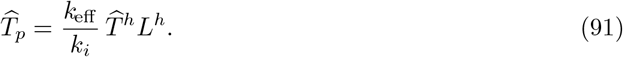

And from the steady-state equation for *dS/dt*,

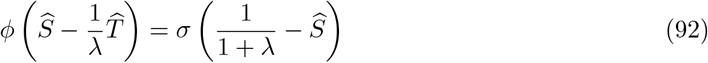

Therefore,

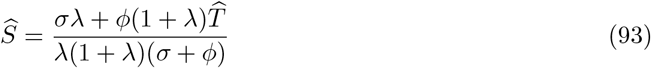

Inserting Eqn. (92) into the steady-state equation for *dT/dt* yields:

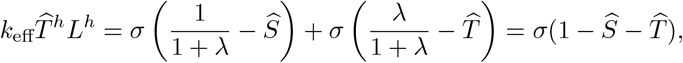

and inserting there Eqn. (93) gives,

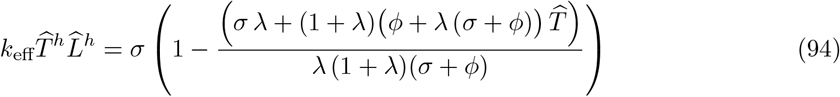

Finally, inserting Eqn. (94) into Eqn. (91) one has:

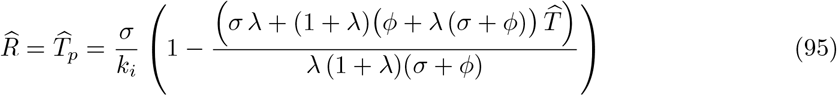

**Step 3:** To calculate *E*_max_ we will proceed in two steps.

First, from Eqn. (94) one has:

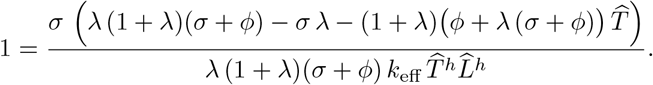

Let’s now assume that 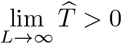. Then, taking the same limit of the above equation one has:

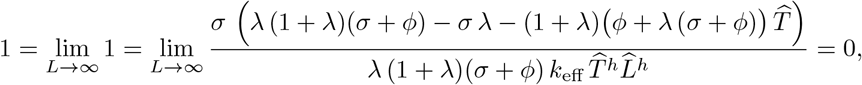

which is a contradiction. Since 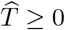, it follows then that 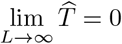.

Second, applying this result to Eqn. (95) we have:

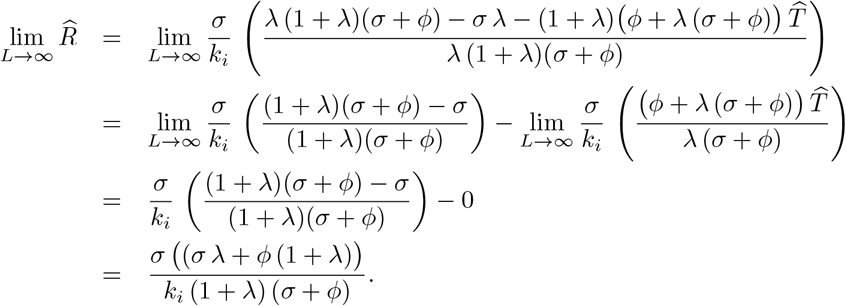

Therefore:

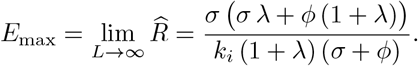

**Step 4:** *EC*_50_ is the value of *L* that satisfies the equation 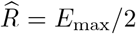. To solve it requires deriving previously 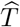 in Eqn. (94) in terms of *L*. However, that equation can be solved analytically only up to *h* = 4. Moreover, after substituting 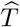 into the generic formula 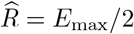, the resulting equation become exceedingly complicated even for *h* = 3. Therefore, in what follows, we present general calculations of *EC*_50_ only for *h* = 1, 2.

**h** = **1**

From Eqn. (94) one has,

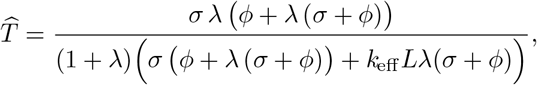

and applying this result to Eqn. (95) yields,

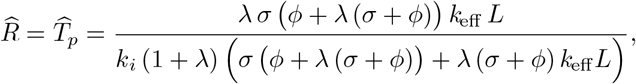

*EC*_50_ is the value of *L* that satisfies the equation:

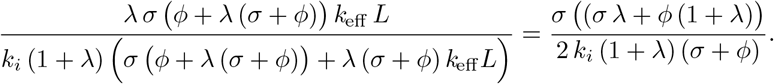

Solving this equation gives:

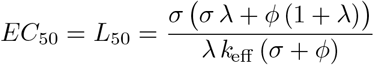

**h** = **2**

From Eqn. (94),

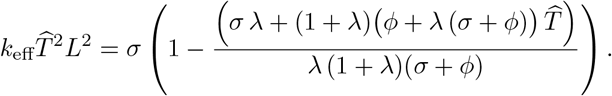

Solving this quadratic equation for *T* gives:

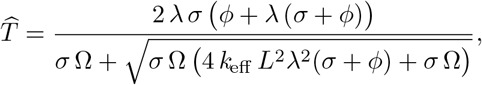

with: Ω = (1 + *λ*)(*ϕ* + *λ*(*σ* + *ϕ*)); and substituting this result into Eqn. (95) yields:

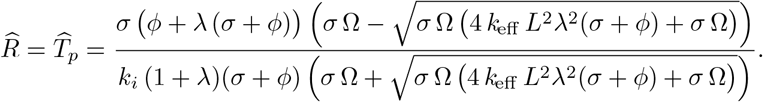

*EC*_50_ is the value of *L* that satisfies the equation:

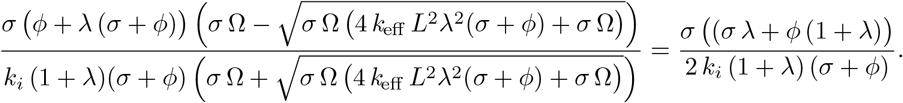

Solving this equation for *L* gives:

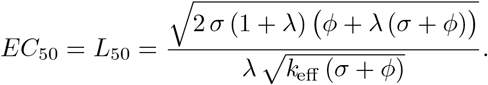

□

